# Consistent cross-modal identification of cortical neurons with coupled autoencoders

**DOI:** 10.1101/2020.06.30.181065

**Authors:** Rohan Gala, Agata Budzillo, Fahimeh Baftizadeh, Jeremy Miller, Nathan Gouwens, Anton Arkhipov, Gabe Murphy, Bosiljka Tasic, Hongkui Zeng, Michael Hawrylycz, Uygar Sümbül

**Author notes:** **Correspondence**: Rohan Gala, Uygar Sümbül.

## Abstract

Consistent identification of neurons in different experimental modalities is a key problem in neuroscience. While methods to perform multimodal measurements in the same set of single neurons have become available, parsing complex relationships across different modalities to uncover neuronal identity is a growing challenge. Here, we present an optimization framework to learn coordinated representations of multimodal data, and apply it to a large multimodal dataset profiling mouse cortical interneurons. Our approach reveals strong alignment between transcriptomic and electrophysiological characterizations, enables accurate cross-modal data prediction, and identifies cell types that are consistent across modalities.

**Highlights:** Coupled autoencoders for multimodal assignment, Analysis of Patch-seq data consisting of more than 3000 cells

## Introduction

The characterization of cell types in the brain is an ongoing challenge in contemporary neuroscience. Describing and analyzing neuronal circuits using cell types can help simplify their complexity and unravel their role in healthy and pathological brain function.^1–6^ This has prompted major consortia such as the BRAIN Initiative Cell Census Network (BICCN) to seek a comprehensive characterization of cell types and their function.^7^ However, the effectiveness of such approaches is predicated on the existence of cellular identities that manifest consistently across different observation modalities, and our ability to identify them. While single cell RNA sequencing (scRNA-seq) experiments have uncovered a detailed transcriptomic organization of cortical cells in the mouse brain,^8, 9^ emerging experimental techniques now enable concurrent characterization of multiple aspects of neuronal identity and function.^7^ For example, MERFISH^10^ can provide paired *in situ* measurement of anatomy and gene expression of multiple neurons in intact tissue, and the Patch-seq protocol^11^ can characterize morphology, electrophysiology, and gene expression of single neurons in tissue slices. Aligning modalities in such paired reference datasets offers the opportunity to move towards a unified, multimodal view of cellular diversity, and potentially enable translation of individual measurements across modalities with high fidelity.

Aligning multimodal data for cell type research is challenging due to complexity of biological relationships between modalities, difficulties in measuring signal and quantifying noise in each modality, and the high dimensional nature of measurements. We present a deep neural network based methodology referred to as *coupled autoencoders* to perform alignment for paired datasets, demonstrate its utility for the multimodal cell type identification problem, and provide an unsupervised, data-driven characterization of GABAergic cell type diversity, which has been a central problem in neurobiology.^5,12–14^ Classical approaches to group GABAergic cells based only on anatomy, physiology, etc. typically disagree on both, the number and the identity of cell types^5^ presumably because the relative importance of the features within an observation modality is unknown. Yet, unequivocal identification of interneurons is essential to elucidate the brain circuits they participate in. Moreover, discordant results cast doubt on the hypothesis that neurons have unique identities, whereby different experiments reveal different facets of those identities, potentially through complicated transformations and noise processes. Here, we focus on the two modalities with the largest number of paired samples in a recent Patch-seq dataset.^14^ There are neither overlapping features nor known associations across the two modalities. Using the fact that the same samples are measured in each modality, our goal is to formulate cell identities which are consistent across these modalities.

## Results

Coupled autoencoders consist of multiple autoencoder networks, each of which consists of encoder and decoder subnetworks. These subnetworks are nonlinear transformations that project input data into a low dimensional representation, and back to the input data space respectively, Figure 1a. In learning these transformations, the goal is to simultaneously maximize reconstruction accuracy for each data modality as well as similarity across representations for the different modalities. In particular, hyper-parameter *λ* (Methods) controls the relative importance of achieving accurate reconstructions versus learning representations that are similar across modalities.

**Figure 1:**
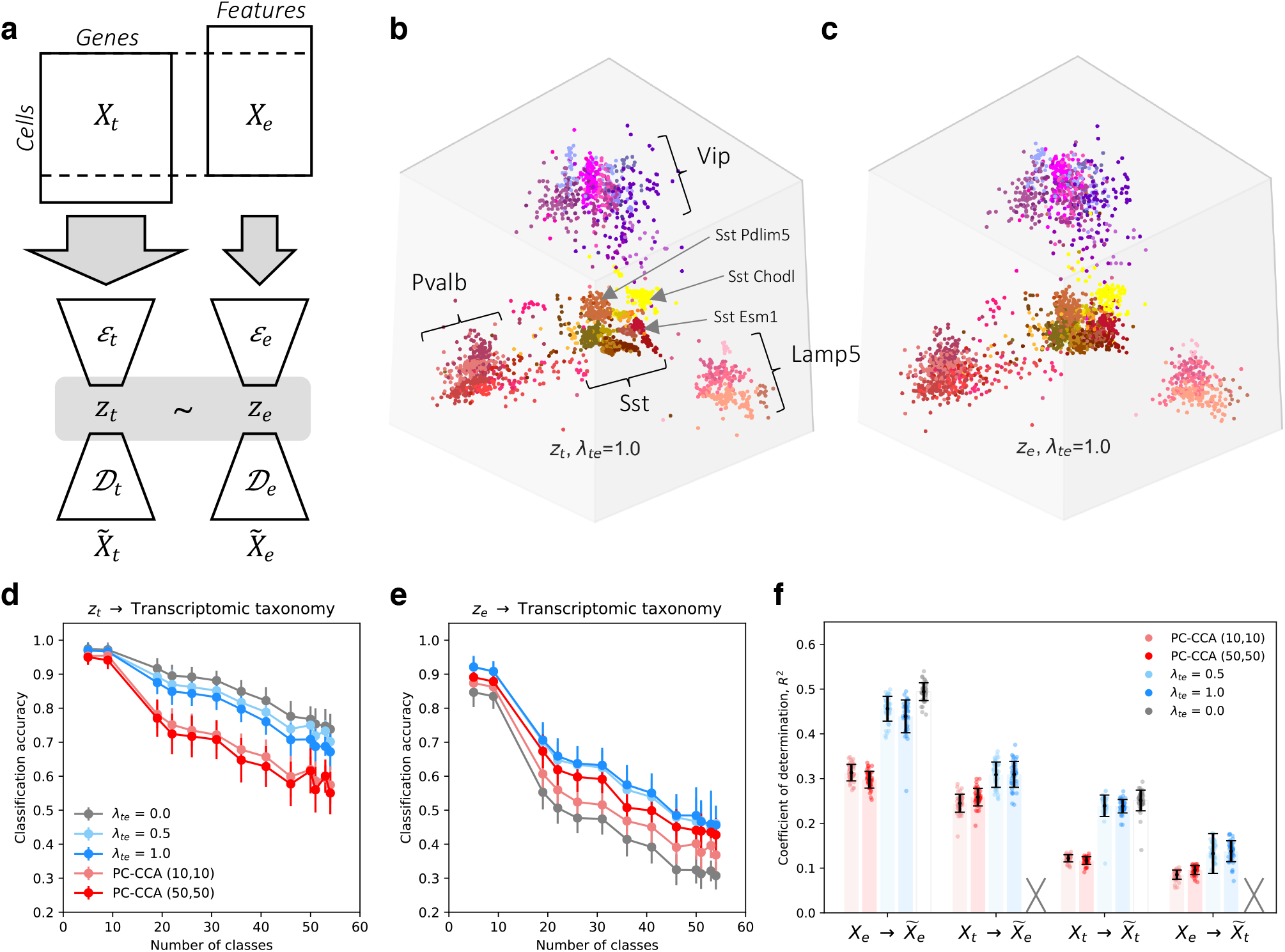
Coordinated representations of transcriptomic and electrophysiological profiles with coupled autoencoders. (a) Schematic showing the coupled autoencoder architecture for Patch-seq data. Encoders (*ε*) compress input data (*X*) into low dimensional representations (*z*). Decoders (*𝒟*) reconstruct data 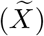 from representations. The coupling penalty in the loss function encourages representations to be similar across the transcriptomic (t) and electrophysiology (e) modalities. (b-c) 3-d coordinated representations of the transcriptomic and electrophysiological datasets. Each point represents a single cell, colored by its cell type membership according to the reference transcriptomic taxonomy. (d-e) Performance on supervised cell type classification tasks, at different resolutions of the reference taxonomy. Classification with QDA is performed using 3-d representations obtained with coupled autoencoders and with linear methods. (f) Performance on within-modality (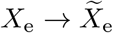 and 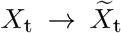), and cross-modality (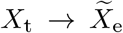 and 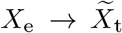) reconstruction tasks. Uncoupled representations are not suitable for cross-modal tasks. Error bars show mean*±*SD over 43-fold cross validation for panels (d-f). Note that there are 1,252 genes vs. 68 electrophysiology features in thedataset, over which (f) is calculated.

If a common latent representation exists across transcriptomic and electrophysiological measurements, which captures salient characteristics of neurons in the individual data modalities, an important consequence will be that unimodal electrophysiological measurements of interneurons can be used to predict gene expression, and vice versa. This ability to translate measurements across modalities may enable researchers to test cell type specific hypotheses without performing costly and potentially intractable multimodal experiments. The ability to align these modalities would strongly support the hypothesis that molecular and electrophysiological properties of individual neurons are closely related, reflecting attributes of a common cell type, albeit through a complicated mapping. Importantly, although linear transformations^15,16^ can align the major cell classes to some extent, a more detailed alignment of features and cell types may require non-linear transformations.

A further consequence of such aligned representations would be the ability to identify, without supervision, cell types of key classes such as GABAergic interneurons in the mouse visual cortex that are consistent across transcriptomic and electrophysiological characterizations of this neuron population. The level of agreement between those clusters and a reference transcriptomic taxonomy of cortical cell types,^8^ and the degree of perturbation of cluster boundaries with respect to that reference taxonomy, can enhance both practical and conceptual aspects of our understanding of cell types.

Aligned 3-d representations *z*_t_ and *z*_e_ for the high-dimensional transcriptomic and electrophysiological profiles *X*_t_ and *X*_e_ obtained with coupled autoencoders are shown in Figure 1b-c. Our experiments suggest that a latent dimensionality of 3 *≤ d ≤* 10 can capture the variability in the dataset, Extended Data Figure 5. Here, we choose *d* = 3 to minimize the number of parameters needed for the downstream unsupervised clustering task. Cells labeled according to the reference transcriptomic taxonomy (Extended Data Figure 1-2) cluster together in representations of both observation modalities. Moreover, the representations largely preserve hierarchical relationships between cell types of the reference taxonomy.

Representations obtained with coupled autoencoders may be used to perform a variety of downstream analyses on complex datasets. We considered supervised classification accuracy in predicting cell type labels at different resolutions (Methods) of the reference taxonomy from *z*_t_ and *z*_e_ in Figures 1d-e, and data reconstruction performance in Figure 1f. First, consider the uncoupled (*λ*_te_ = 0.0) setting, where each autoencoder performs nonlinear dimensionality reduction independently for its respective input modality. Representations based on the transcriptomic data alone (*z*_t_, *λ*_te_ = 0) are best suited for supervised cell type classification using quadratic discriminant analysis (QDA), leading to 0.74 *±* 0.05 accuracy for leaf node cell type labels, Figure 1d. This is not surprising, since the reference transcriptomic taxonomy was derived from analyses of gene expression alone. Electrophysiological profiles are expected to be noisy, and of lower resolution compared to transcriptomic profiles.^17^ Nevertheless in Figure 1e, classifiers based on representations of electrophysiology alone (*z*_t_, *λ*_te_ = 0) predict leaf node cell type labels with an accuracy of 0.31 *±* 0.04 (chance level is 0.03). To add context, a model trained only to classify leaf node cell types based on electrophysiological profiles alone led to an accuracy of 0.52 *±* 0.04 (Methods). Lastly, within-modality reconstruction accuracies of uncoupled representations provide an upper limit for both, within- and cross-modal reconstructions that may be achieved with 3-d representations obtained with coupled autoencoders.

To evaluate whether complicated, non-linear transformations underlie the relationship between the transcriptomic and electrophysiological features of neurons, we considered the performance of linear methods (PC-CCA), and coupled autoencoders with *λ*_te_ *∈* {0.5, 1.0} at these tasks, with the representation dimensionality set to 3. The weak dependence on *λ*_te_ in Figure 1 and Extended Data Figure 3 suggests that our method is robust with respect to this hyperparameter. We note that the Patch-seq experiment provides, to the extent of experimental measurement, perfect knowledge of *anchors* between the modalities by virtue of paired measurements. In this setting, the popular tool Seurat^18^ uses a variant of linear CCA to achieve alignment, for which the performance is expected to be comparable to baselines considered here. Results in Figure 1d-f show that coupled autoencoders learn well-aligned representations of transcriptomic and electrophysiology data, such that cell type labels can be predicted with better accuracy, and the cross-modal data can be predicted more reliably compared to linear methods. Importantly, the within-modality reconstruction error is comparable to that obtained in the uncoupled setting, demonstrating that coupled representations enable alignment across modalities while faithfully compressing the individual data modalities.

Cross-modal data prediction (Extended Data Figures 6-7) is a key computational tool for identifying corresponding properties of cell types, and guiding the design of new experiments. We considered a subset of genes that underlie recently discovered cell type specific paracrine signaling pathways in the cortex.^19^ The Patch-seq transcriptomic data shows these cell type specific gene expression patterns, Figure 2a. We used only electrophysiology features to infer the expression patterns for all genes in the cross-modal setting, and show results for the same subset of genes as before. The striking similarity of these expression patterns (Pearson’s *r* = 0.89*±*0.10, mean*±*SD over cell types, Figures 2b and Extended Data Figure 6) demonstrates the effectiveness of coupled autoencoders at the cross-modal prediction task at a granular level. Similar results were obtained for GABAergic cell type marker genes, Supplementary Figure 1. Neuropeptide precursor genes and their cognate G-protein coupled receptors have widespread expression in the cortex and are implicated to form cell type specific broadcast communication networks.^19,20^ Therefore, the high degree of similarity in Figure 2a,b provides indirect evidence for coordination between intrinsic cellular electrophysiology and circuit-level neurotransmitter networks in the cortex. (A link between electrophysiology and cell adhesion molecules was previously studied using scRNA-seq.^21^)

**Figure 2:**
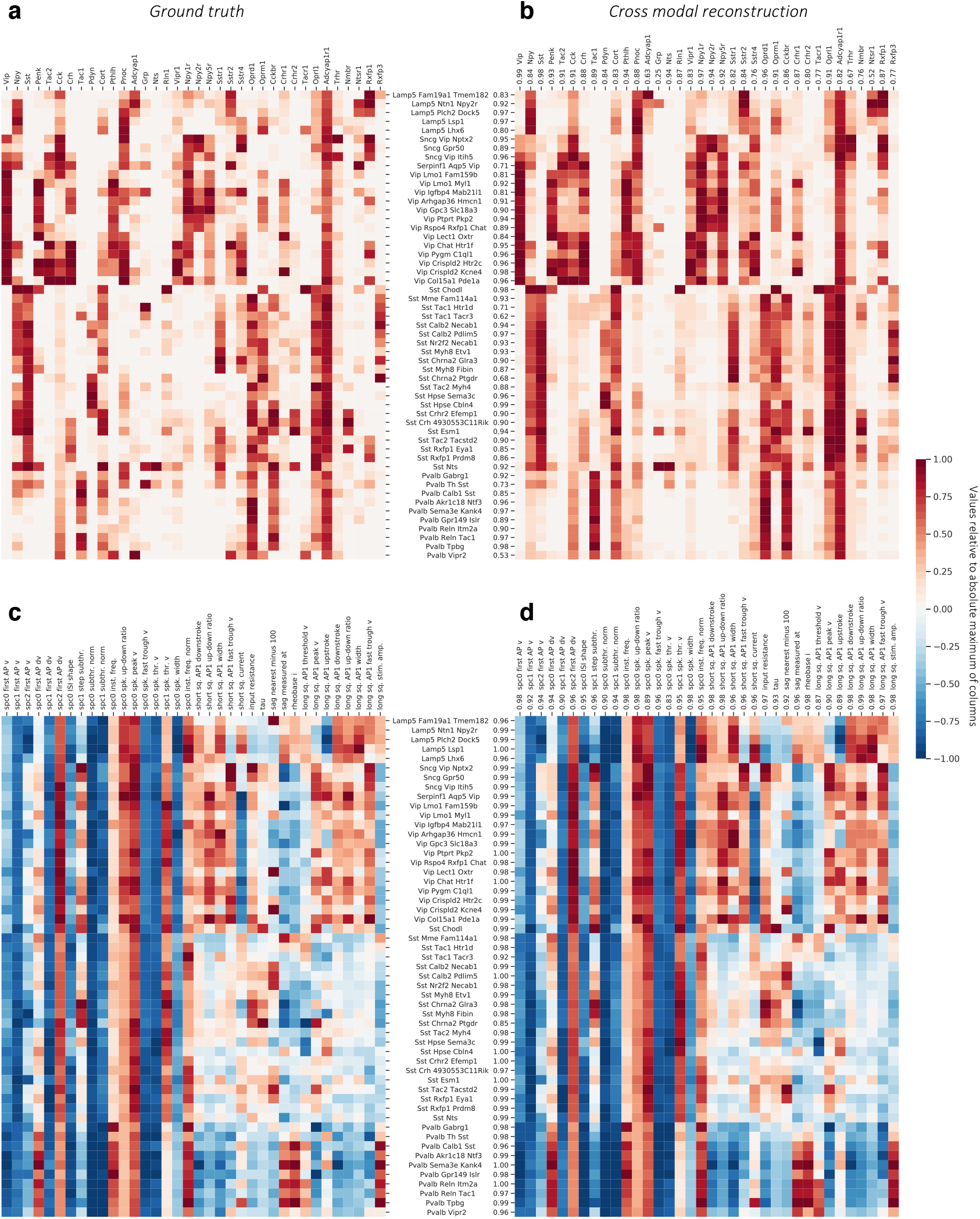
Cross-modal reconstructions capture cell type specific gene expression patterns and electrophysiological features. (a) Gene expression levels averaged over samples reference taxonomy cell types, normalized per gene by the maximum value of each column. (b) Cell type specificity of different genes is captured well by cross-modal prediction of gene expression profiles from electrophysiological features (Pearson’s *r* = 0.89 *±* 0.10, mean*±*SD over cell types). (c) A subset of electrophysiological features pooled by cell types shows analogous cell type specificity. (d) Cross-modal reconstructions of the electrophysiology features from gene expression profiles match the measured electrophysiology features (Pearson’s *r* = 0.98 *±* 0.02, mean*±*SD over cell types). Cell type- and feature-wise fidelity of reconstructions are quantified with Pearson’s *r* for each row and column in (b) and (d) as compared to ground truth in (a) and (c).

We considered cross-modal prediction of electrophysiological features in an analogous manner, pooling values of the features on a per cell type basis, and focusing on features that are captured by the compressed representation well (within-modality reconstruction *R*^2^ *>* 0.42, Extended Data Figure 7). While results of Figure 1d-e already suggest that the electrophysiology features are not as specific to transcriptomic cell types, we can nevertheless identify cell type specific patterns, Figure 2c. The cross-modal reconstruction of these features also matches the data (Pearson’s *r*=0.98*±*0.02, mean*±*SD over cell types), reinforcing the result that gene expression can explain many intrinsic electrophysiological features accurately, and that coupled autoencoders are a powerful starting point to unravel such non-linear relationships. Moreover, the per-feature prediction accuracy can help uncover the features that are important for neuronal identity. (e.g., *Vip, Sst, Npy1r*, and *Oprd1*, up-down ratio of action potential, rheobase current, Extended Data Figures 6-7)

We directly tested the idea that pre-trained coupled autoencoders can be used to predict unobserved cross-modal features in independent experiments by using two recent Patch-seq datasets,^22,23^ which include 107 and 524 inhibitory neurons from mouse motor cortex respectively. We applied a coupled autoencoder trained on the dataset in this work without additional training to predict the transcriptomic labels and electrophysiological properties of these neurons from their transcriptomic profiles. Results in Supplementary Figure 5-6, Extended Data Figures 9,10 show that this approach yields accurate prediction of cell type labels and certain electrophysiological properties, despite a 5% mismatch between the gene lists, differences in electrophysiology protocols and brain regions.

While clustering of individual modalities into cell type taxonomies shows general correspondence, a strategy for consensus clustering is less clear. The notion of a finite set of cell types can be formalized as a statistical mixture model, whereby the observation for each cell is explained by a combination of its membership to one of a discrete number of types, and continuous variability around the type representative. We explored the extent to which such a model can explain the data consistently across modalities. We performed unsupervised clustering on coordinated representations obtained with the coupled autoencoder to explain both modalities. Figure 3a shows the distribution (32.19 *±* 3.16, mean*±*SD) of optimal number of Gaussian mixture components over representations obtained with different network initializations. We take the ceiling of the mean (33) as the number of clusters that can be consistently defined with coordinated representations, and refer to this *de novo* clustering as consensus clusters. Figures 3b and Supplementary Figure 4 demonstrate that the same consensus cluster can be assigned to neurons with high frequency, based on observing either the transcriptomic or electrophysiological (but not both) modality. While the dominant diagonal of this contingency matrix indicates the success of this notion of consistent, multimodal cortical cell types, the off-diagonal entries point to imperfections of this view, either due to experimental noise and limitations of experimental characterization, or due to imperfection of the model itself. As a metric of the consensus between assignments across modalities, we calculated the ratio of clusters for which the diagonal entry of the contingency matrix is at least as large as the off-diagonals of the corresponding row and column. For the reference labels, we obtained *c*_ref_ = 0.26 *±* 0.01, while for the consensus clusters, *c*_con_ = 0.87 *±* 0.04 on all cells. For test cells, we obtained *c*_ref_ = 0.26 *±* 0.01 and *c*_con_ = 0.58 *±* 0.03 (Methods, Supplementary Figures 3-4). These results suggest that consensus clusters can be used to produce cell type assignments for which there is a better agreement across experimental modalities, compared to the reference taxonomy.

**Figure 3:**
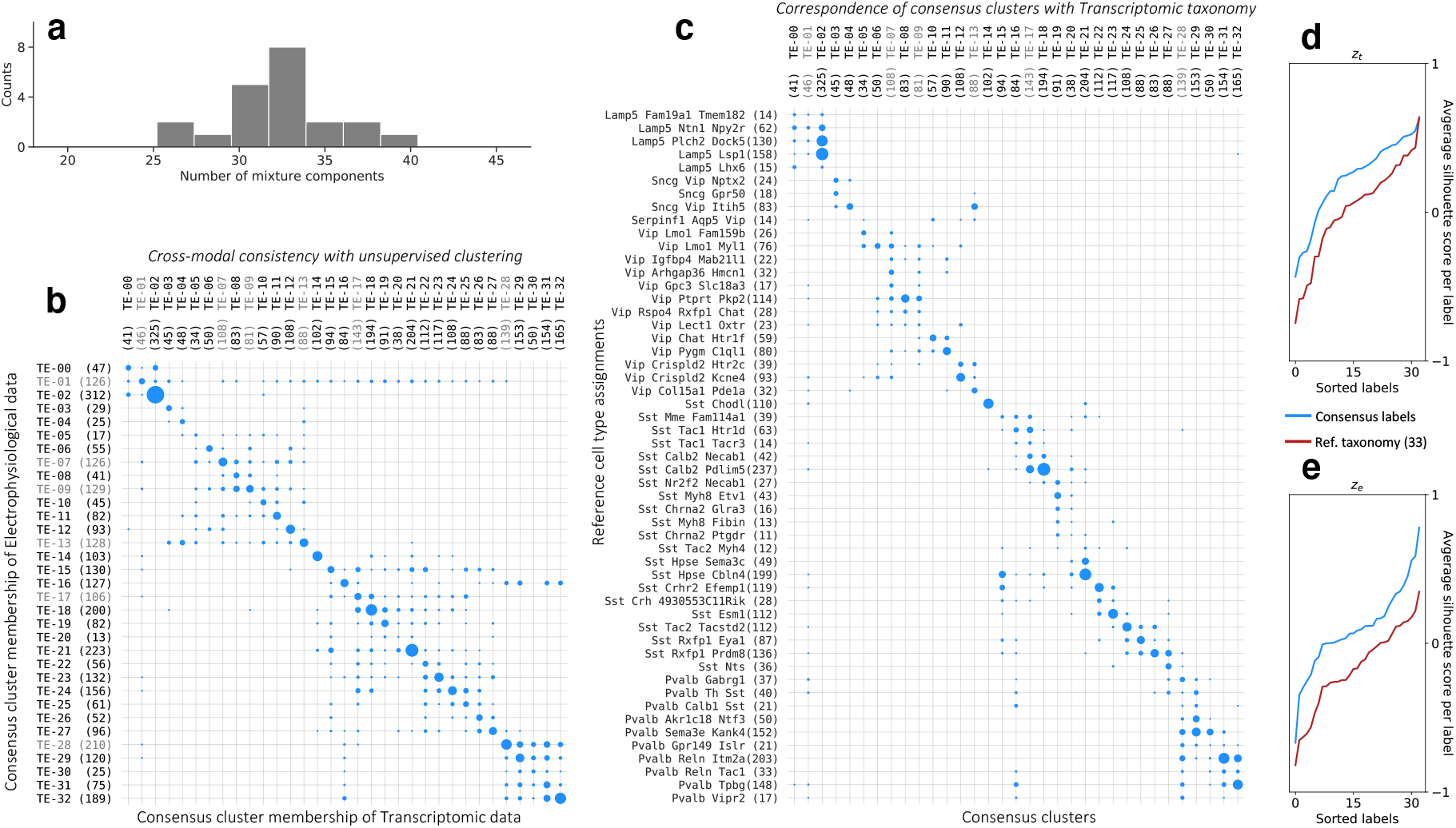
Deriving a consensus cell type clustering. (a) Unsupervised clustering using Gaussian Mixtures on the coordinated representation *z*_t_ and BIC based model selection suggests 33.0 consensus clusters (32.19*±*3.16, mean*±*SD). (b) Contingency matrix for cluster assignments based on independent, unsupervised clustering of the transcriptomic and electrophysiology representations shows that the clusters are highly consistent. (c) Contingency matrix for the leaf node cell type labels of the reference hierarchy compared to unsupervised cluster assignments show that these unsupervised clusters have substantial overlap with known transcriptomic cell types. Number of cells for each label are indicated within parentheses next to the label, and area of the dots is proportional to number cells in the scatter plots of (b-c). (d) Consensus clusters are compared with an equal number of cell classes obtained by merging the reference hierarchical taxonomy, using silhouette analysis based on coupled representations. Average per-cluster silhouette values for test cells are larger for consensus cluster labels. Clusters for which the silhouette score is less than 0 in (d) and (e) are grayed out in panels (b-c).

The consensus clusters are also consistent with the reference taxonomy, although not to the degree of all leaf node labels, Figure 3c. This can indicate over-split (e.g., Lamp5 Plch2 Dock5, Lamp5 Lsp1), under-split (e.g., Sst Calb2 Pdlim5), and not-tight (e.g., Vip Lmo1 Myl1, Sst Mme Fam114a1) cell types in the reference taxonomy. To quantify the degree to which different label sets represent the underlying data, we report the average silhouette score for test samples (not used to train the coupled autoencoder or the mixture model) for each label, using *z*_t_ and *z*_e_, and compare consensus clusters against those of the reference transcriptomic taxonomy (Figure cd,e). A negative silhouette score suggests an unreliable cluster (greyed out in Figure 3b,c). Not only do consensus clusters capture the structure of the data better than the reference labels on *z*_t_ (average silhouette score for consensus clusters *s*_con_ = 0.24 *±* 0.01 and for reference labels *s*_ref_ = 0.14 *±* 0.01, mean*±*SD over 5 best initializations), but also on *z*_e_ (*s*_con_ = 0.04 *±* 0.02, *s*_ref_ = *−*0.13 *±* 0.01). For the reference taxonomy, we repeated this analysis using stand-alone (rather than aligned) representations and for different hierarchical mergings of the taxonomy with at least 33 labels, and we obtained very similar results. (Methods, Extended Data Figure 8). These results suggest that consensus clusters are a more identifiable characterization of the joint transcriptomic and electrophysiological diversity of interneurons than one based on transcriptomics alone.

## Discussion

Our analysis of the largest multimodal Patch-seq dataset to date with unsupervised clustering on coordinated representations reveals *∼*33 clusters that can be defined consistently with transcriptomic and electrophysiological measurements of cortical GABAergic neurons, providing a deeper association of these modalities than previously explained. Beyond inferring cell types, coupled autoencoders trained on reference datasets can serve as efficient “translators” for experiments using a single observation modality to infer neuronal properties in other modalities. This capability can provide indirect evidence for/against hypotheses that are hard to test, such as predicting the expression levels of genes regulating ion channels of interest, purely from observations of intrinsic electrophysiology.

An intriguing and essential issue regarding cell types is to what extent they are inherently discrete entities or representatives of a continuum.^24^ A mixture model allows for types to overlap each other in the representation space so long as the cluster centers are more dominant than the peripheries. With this model, mouse visual cortex interneuron Patch-seq data suggests the existence of 33 clusters, more than the *∼*5 well-known subclasses but less than the *>* 50 partitions suggested by scRNA-seq data alone. A potential caveat is that scRNA-seq in the Patch-seq experiment is noisier than its stand-alone counterpart.^14^

Coupled autoencoders can jointly analyze multiple modalities. Future work can incorporate additional observation modalities (e.g. morphology, connectivity) to improve our understanding of neuronal identity.

Finally, dataset size plays an important role in all our results. More samples can allow the use of larger representation space dimensionality and improve cross-modal data prediction. Similarly, clustering is ill-defined for cell types with too few samples; further analysis of consensus or transcriptomic clusters (Figure 3c) should take sample size into account. Therefore, the number of cortical GABAergic interneuron types is likely to grow, and the number of consensus clusters in Figure 3 more likely represents an under-count of the diversity when the notion of cell types is considered as a mixture model.

## Methods

### Coupled autoencoders

Approaches to discover and extract relationships in multimodal datasets are discussed in literature as cross-modal retrieval, multimodal alignment, multi-view representation learning.^25–27^ Deep learning methods such as DeepCCA^28,29^ and correspondence autoencoders^30^ are promising approaches to achieve multimodal data alignment, but have had limited success in associating complex neural datasets (See Supplementary Information for an overview of modern data alignment approaches). Our coupled autoencoder networks are related architectures with key improvements to scaling of representations that are critical for the overall quality of learned representations.^31^

We first describe the general coupled autoencoder framework. Then, we show its application to the Patch-seq dataset. For *K* observation modalities, we represent the coupled autoencoder by

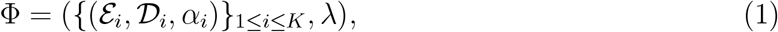

where *ε*_*i*_ and 𝒟_*i*_ denote the encoding and decoding networks for the *i*-th observation modality, *α*_*i*_ sets the relative importance of the different modalities, and *λ ≥* 0 sets the relative importance of representation fidelity within observation modalities versus the alignment of different representations.

For a set of paired observations *X* = {(*x*_*s*1_, *x*_*s*2_, …, *x*_*sK*_), *s ∈ S*}, we define the loss due to Φ as

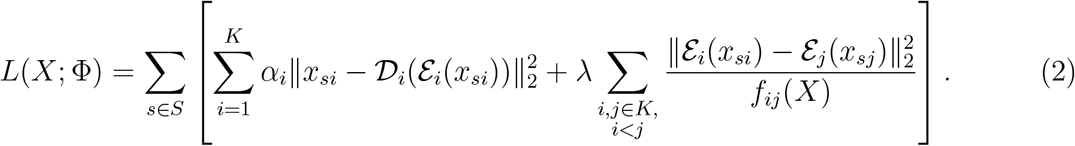

That is, each autoencoding agent (Figure 1a) within the coupled architecture processes a separate data modality and optimizes a loss function that consists of penalties for (1) the discrepancies between the actual input *X* and reconstructed input 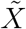 (2) mismatches between the representations learned by the different agents. (A slightly more general treatment can be found in Ref.^31^)

In Eq. 2, the functional form of the denominator *f*_*ij*_ that scales the mean squared difference between representations of the same sample based on the different data modalities, is crucial to learn good quality representations. Common choices for *f*_*ij*_ lead to pathological solutions, i.e. the latent representations collapses into a zero- or one-dimensional space (see Propositions below). To avoid such pathological solutions, we propose using:

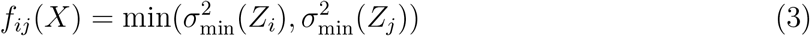

where *σ*_min_(*Z*_*i*_) denotes the minimum singular value of the matrix *Z*_*i*_, which consists of rows *Z*_*i*_(*s*, :) = *z*_*si*_ where *z*_*si*_ = *ε*_*i*_(*x*_*si*_). In practice, we perform stochastic gradient descent and calculate *f*_*ij*_ by its mini-batch approximation. Scaling the coupling loss term in this manner approximates whitening by the full covariance matrix well, and also is practically important when the batch size is small or representation dimensionality is large, regimes where calculating the full covariance matrix would be unreliable and computationally expensive.

### Representations collapse for common scaling function choices

Proofs for the following propositions can be found in Ref.^31^

**Proposition 1**. *Let f*_*ij*_ = 1. *Representations of the coupled autoencoder that minimize the loss in Eq. 2 satisfy* ||*z*_*si*_||*< ϵ, for any norm* || *·*||, *input set X, ϵ >* 0, *and all s, i*.

**Proposition 2**. *Let f*_*ij*_ *implement* Batch Normalization^32^. *Representations of the coupled autoencoder that minimize the loss in Eq. 2 satisfy* 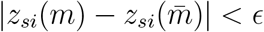, *for all* 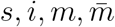, *and ϵ >* 0, *with probability* 1.

### Application to the Patch-seq dataset

We use the fact that the same neurons were profiled with both modalities to obtain aligned, low-dimensional representations of the gene expression profiles and electrophysiological features. In the case of just these two data modalities, transcriptomics (*t*) and electrophysiology (*e*), the loss function according to Eq. 2 consists of two reconstruction error terms, and a single coupling error term. For a single sample *s*,

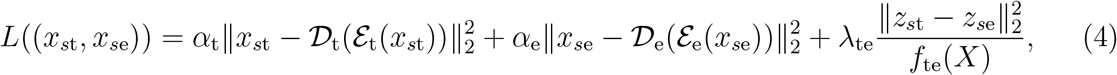

where *z*_*s*t_ = *ε*_t_(*x*_*s*t_) and *z*_*s*e_ = *ε*_e_(*x*_*s*e_). Here *x*_*s*t_ denotes gene expression vector for sample *s* and *x*_*s*e_ denotes the concatenated sPC and physiological feature measurement vectors for the same sample. The interplay between the accuracy with which the representations capture the individual data modality, versus how well the representations are aligned is a fundamental trade-off that any attempt to define consistent multimodal cell types must resolve (see Supplementary Information for an equivalent formulation in the probabilistic setting). The hyper-parameters *α*_t_, *α*_e_ and *λ*_te_ explicitly control this trade-off in our formulation (Extended Data Figure 3 shows behavior over a range of these values with the Patch-seq dataset). We set all three parameters to 1.0 for all central analyses in this manuscript.

### Data augmentation

Data augmentation is important to regularize the networks and alleviate overfitting, particularly when the dataset size is small. We mimicked the biological dropout phenomenon^33^ and used Bernoulli noise (i.e., Dropout^34^) to augment repeated presentations of the transcriptomic vectors while training. This strategy also renders the network robust to partial mismatches in gene lists, and reduces dependence of the representations and reconstructions on specific marker genes. The individual electrophysiological features have unequal variances, since the total variance in the sPC is normalized on a per-experiment basis. We therefore used additive Gaussian noise with variance proportional to that of the individual features to augment the electrophysiological vectors while training the network.

The reconstruction loss for the decoders was calculated with both, the representation obtained by the encoder network of the same modality, and that obtained by the encoder for the other modality. This was done to improve performance of cross-modal prediction. This way of calculating the reconstruction loss can be viewed as an augmentation strategy for the decoder networks, that significantly improves the accuracy of cross-modal prediction (Extended Data Figure 4).

### Linear baselines

Canonical correlation analysis (CCA) is a standard linear method to align low dimensional representations.^15^ To optimize the performance with linear methods, we first used principle component analysis (PCA) to reduce the dimensionality of individual data modalities, followed by CCA to achieve aligned representations across the modalities. The number of dimensions to which the transcriptomic and electrophysiology data were reduced to with PCA is indicated as a tuple in the legends of Figure 1. The dimensionality of CCA representations was chosen to match the dimensionality obtained with coupled autoencoders (dim=3). The inverse CCA and PCA transformations were used to reconstruct data from the representations both, for the the within- and across-modality cases in Figure 1f. Reconstruction performance for different representation dimensionality is compared in Extended Data Figure 5.

### Supervised cell type classification

Label sets obtained at different resolutions of the reference transcriptomic taxonomy were used as ground truth to evaluate representations. The different resolutions correspond to different horizontal levels of the reference taxonomy hierarchy in Extended Data Figure 1. Starting from the leaf node cell type labels, each cell is assigned the parent node label based on the set of labels that remains at a given level of the hierarchy. Thus at the lowest resolution, there is just a single class (n59) encompassing all neurons, and at the highest resolution there are 53 classes i.e. cell type labels (Extended Data Figure 1). Quadratic Discriminant Analysis (QDA)^15^ was used to perform classification with the representations obtained with coupled autoencoders or CCA, and used to predict the cell type labels for all such label sets. Cells that were not used to train the coupled autoencoder were used to obtain accuracy values shown in Figure 1(d-e) using a *k*=43 fold cross validation approach. Validation folds were obtained such the class distribution in each fold was similar to that for the overall dataset. Classes with *n ≤* 10 samples in the dataset were discarded from the analysis. Similarly, classes for which there were less than *n*=6 samples in the training set of any fold were discarded from evaluation for only that fold, since QDA classifier parameters for those poorly represented classes would be unreliable. The results were pooled across the folds for the remaining number of classes (i.e. QDA components) in Figure 1(d-e). The architecture of the neural network only trained to classify cell types at the highest resolution of the taxonomy using only electrophysiological profiles used the same encoder network as for the *X*_e_ autoencoder, except that the output was a 53 way classification. The network was trained with the standard cross entropy loss for classification.

### Unsupervised clustering and consensus clusters

For this analysis, 80% of the cells were used for training, and the remaining 20% served as the test set. The training and test sets had similar distributions of the cell type labels based on the reference taxonomy. The coupled autoencoder (*λ*_te_ = 1.0) was initialized 21 times to obtain as many different representations. For each representation, Gaussian mixture models with different numbers of components (15 to 45 in steps of 1) were fit on *z*_t_ utilizing only the training set. The model with the lowest value of the Bayesian Information Criterion^15^ on the training set was used to determine the optimal number of components. The distribution for optimal number of mixture components across the 21 different representations was binned using the Freedman-Diaconis rule,^35^ Figure 3a. Based on this distribution (32.19 *±* 3.16, mean*±*SD) we use the ceiling of the mean as the number of clusters that can be consistently defined with coordinated representations. These 33 mixture components are referred to as consensus clusters. To show results on the test cells, we picked the representation from coupled autoencoder model with the best total reconstruction error. The 33 component Gaussian mixture model was then used to assign consensus cluster labels to test cells based on *z*_t_, as well as based on *z*_e_. The consensus cluster assignments obtained in this manner are compared in Figure 3b. (Supplementary Figure 2 shows the individual BIC plots for the different representations, and offers a comparison with a similar analysis using PC-CCA representations) We used the Hungarian algorithm to match the consensus clusters with leaf node cell types of the reference taxonomy, using the negative of the contingency matrix based on training cells as the cost function. The order of the consensus clusters in Figure 3b-c reflects this optimal match.

### Evaluating the agreement and consistency of cluster assignments

To evaluate the agreement between assignments across the experimental modalities on a per cluster basis, we calculated the following fraction:

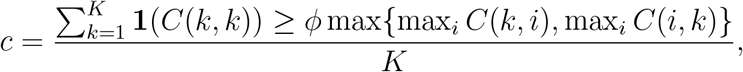

where *K* denotes the number of clusters, *C* denotes the contingency matrix for which the fraction is calculated, and ***1*** denotes the indicator function; ***1***(*a*) = 1 if *a* holds, otherwise ***1***(*a*) = 0. *c*_con_ (*c*_ref_) refers to this quantity when calculated with the contingency matrix calculated over the consensus labels (reference taxonomy merged to 33 labels). We set the scalar factor *ϕ ≥* 1 to *ϕ* = 1 in the reported experiments. We also found that the conclusion that *c*_con_ *> c*_ref_ is not sensitive to this factor over a broad range of values (1 to 5). To obtain uncertainty estimates, we selected the best 5 of the 21 networks used for training on the same 80% of the dataset with different initializations, based on the lowest total reconstruction error on the test set. We performed clustering with 33 mixture components to obtain the labels and calculate the corresponding contingency matrices. For any other label set, we use the representation to train QDA classifiers and assign labels to the test set to obtain the contingency matrix.

To evaluate the ability of a set of labels in representing the underlying data, we used the Silhouette analysis. While the Silhouette values reported in the main text, *s*_con_ and *s*_ref_, are averages over all involved samples, the values reported in the corresponding figures represent averages on a per cluster basis. To obtain uncertainty estimates, we selected the best 5 of the 21 networks used for training on the same 80% of the dataset with different initializations, based on the lowest total reconstruction error on the test set. Silhouette scores were obtained for the different label sets using the test set representations obtained from these networks.

### Patch-seq dataset

We used the transcriptomic and electrophysiological profiles of GABAergic interneurons from mouse visual cortex of a recent Patch-seq dataset.^14^ Briefly, neurons were patched with biocytin-filled electrodes with which the electrophysiological responses to a series of hyperpolarizing and depolarizing current injections were recorded, nuclear and cytosolic mRNA was extracted, reverse transcribed and the resulting cDNA was sequenced using SMART-Seq v4(Tasic et al., 2018). Among the 3,708 cortical GABAergic interneurons reported in that work, axonal and dedritic morphology of only 350 cells was reconstructed. We restricted our analysis to the two modalities with largest number of samples, and dropped the morphology modality altogether. The 3,708 cells were also mapped to the reference transcriptomic hierarchical taxonomy^8^ with different levels of confidence. We discarded cells which were annotated as “inconsistent”^14^ based on this confidence level, leaving 3,411 cells for which both transcriptomic and electrophysiological profiles were available. The relevant taxonomy, and abundances of cells per type for well-representated types (at least 10 samples per type) are shown in Extended Data Figures 1 and 2.

A set of 1,252 genes used as input for the analyses in this study. The gene selection procedure included two filtering steps. The first step excluded genes where the primary source of variation in gene expression is unlikely to be related to its cell type identity. Specifically, we removed (1) genes that are highly expressed in non-neuronal cells, (2) genes with reported sex or mitochondrial associations, and (3) genes that are much more highly expressed in Patch-seq data vs. Fluorescence Activated Cell Sorting (FACS) data (or vice versa) and therefore may be associated with the experimental platform^14^. We also removed gene models and some other families of unannotated genes that may be difficult to interpret. The second filtering step used the *β* score, which is a published measure of how binary a gene is with respect to cell types.^36^ Higher *β* indicates that for each cell type a gene is either expressed (with CPM>1) or unexpressed in most cells. We removed all genes with *β <* 0.4, leaving a total of 1,252 genes in the analysis after both filtering steps. Gene expression values were CPM normalized, and then log_*e*_(· + 1) transformed.

44 sparse principle components (sPC) were extracted to summarize the time series data from different portions of the electrophysiology measurement protocol.^14^ Additionally 24 measurements of intrinsic physiology features were obtained using the IPFX library https://ipfx.readthedocs.io/. The sPC values were scaled to have unit variance per experiment. The remaining features were individually normalized to have zero mean and unit norm. Note that the coupled autoencoder trains to minimize the mean squared error for both modalities. Therefore, normalization of individual features on the electrophysiology side can affect the *R*^2^ calculations (the autoencoder may also act as a denoiser and impute genes that suffer from experimental drop-out which can affect the observed *R*^2^ values as well.) Data was divided into *k*=43 folds for cross validation experiments. For the consensus cluster experiments, 20% of the cells were set aside as the test set. Different random seeds were used to train networks 21 times on the remaining 80% of the cells.

### Cross modal reconstruction of inhibitory cell type marker genes

We compiled a set of marker genes for inhibitory cell types from (according to Figures 5e and 5f in ^8^) to showcase the cross-modal reconstruction ability of our model. Out of these, 38 genes were available in the set of 1,252 genes used as input and cross-modal reconstructions for these are shown in Supplementary Figure 1. Note that 6 of these marker genes are also present in the list of neuropeptide precursor gene list shown in Figure 2.

### Supervised approach to identify consistent cell types based on reference taxonomy

Our experiments suggest that one can identify *∼* 33 clusters with in this dataset using a completely unsupervised approach. To argue that clusters obtained in this manner are more consistent across modalities, we compare the contingency matrix of Figure 3b with one obtained with a supervised approach, utilizing the labels of the reference taxonomy, Supplementary Figure 3. Accordingly, we first merged the reference hierarchy of Figure 1 to obtain 33 class labels. With these labels, we trained separate QDA classifiers on 3 dimensional representations *z*_*t*_ and *z*_*e*_ from the uncoupled (*λ*_*te*_ = 0) autoencoders. Note that this is equivalent to performing dimensionality reduction with separate autoencoders trained on the individual modalities. First, we show a representative contingency matrix for the ground truth transcriptomic labels, and the labels predicted by the classifier for test cells. The dominant diagonal in Suppplementary Figure 3a shows that the uncoupled 3 dimensional representations capture the transcriptomic classification very well. Suppplementary Figure 3b shows the contingency matrix for label predictions with *z*_*t*_ and *z*_*e*_. We observe that for certain classes, classifiers trained on different modalities never lead to identical labels (zeros along the diagonal). We quantify this by defining a fraction of co-occupied labels, *f*_*co*_. For Figure 3 *f*_*co*_ = 0.88, while for the supervised approach devised here, *f*_*co*_ = 0.42 *±* 0.02 (mean*±*SD, 6-fold cross validation).

### Application as a cross-modality translator for unimodal data

We used two different published Patch-seq datasets with transcriptomic and electrophysiological profiles of GABAergic cells, Scala et al. 2019^22^ and Scala et al. 2020,^23^ to demonstrate the utility of coupled autoencoders to serve as translators for unimodal data. The Patch-seq data used in the main text is used as the reference dataset, and is referred to as the Gouwens et al. 2020 dataset. The Scala et al. 2019 dataset consists of 107 neurons, and Scala et al. 2020 dataset consists of 524 neurons sampled in mouse motor cortex.

Previous studies have shown that the cell type diversity of inhibitory neurons is essentially conserved across brain areas.^8^ Therefore, even though both, the Scala et al. 2019 and the Scala et al. 2020 datasets profile neurons from mouse motor cortex, we can hope to use the mouse visual cortex Patch-seq dataset used in this study to serve as a meaningful reference. Nearly 5% of the genes that were used as input for the coupled autoencoders were either not measured, or were missing from the transcriptomic profiles of neurons in both these datasets. At the time of network training, we zeroed out the genes expression values at random both to mimic gene dropout^33,34^ and to increase the robustness against non-identical input gene lists. Therefore we were able to use the pre-trained network without additional training to make predictions with these datasets.

The Scala et al. 2020 dataset was obtained from the public repository related to this work, https://github.com/berenslab/mini-atlas commit #1a0be. The dataset consists of gene expression profiles that were mapped to the reference taxonomy considered here. We relied on this mapping to select 524 cells that were confidently mapped to the inhibitory types that were well sampled in the Gouwens et al. dataset. In particular, we filtered out cells that were mapped to a single leaf node of the reference taxonomy with less than 80% confidence.

Results for inferring transcriptomic cell types and predicting electrophysiological features from gene expression with pre-trained coupled autoencoders are shown in Supplementary Figures 5-6, Extended Data Figures 9,10.

The electrophysiology measurement protocols in both these datasets differ from the one in the Allen Institute dataset. In particular, differences in the temperature and internal/external solutions with which experiments were conducted are expected to contribute to differences in estimated parameters across the datasets.

## Code availability

Code for the coupled autoencoder implementation and analysis are available at https://github.com/AllenInstitute/coupledAE-patchseq. An interactive version of the code base is provided in the Code Ocean capsule https://doi.org/10.24433/CO.4098627.v2. ^37^

## Data availability

The Patch-seq transcriptomic data is available at http://data.nemoarchive.org/other/grant/AIBS_patchseq/transcriptome/scell/SMARTseq/processed/analysis/20200611/, and the electrophysiological data is available at https://dandiarchive.org/dandiset/000020. For the Scala et al. 2019 dataset, the sequencing data is available under accession number GSE134378, and electrophysiological data is available at https://doi.org/10.5281/zenodo.3336165. The Scala et al. 2020 dataset was obtained from the public repository related to this work, https://github.com/berenslab/mini-atlas commit #1a0be.

## Acknowledgements

We wish to thank the Allen Institute for Brain Science founder, Paul G Allen, for his vision, encouragement and support. This work was supported by the NIH grant 1RF1MH123220-01.

## Contributions

R.G., U.S.: Methodology, Investigation, Writing – original draft and revision. R.G.: Software, Formal analysis. A.B., F.B., J.M., N.G., A.A., G.M.: Data curation and pre-processing. B.T., M.H.: Writing – revisions. R.G., A.B., F.B., J.M., N.G., G.M., B.T., H.Z., M.H., U.S.: Conceptualization.

## Extended Data

**Extended Data Figure 1:**
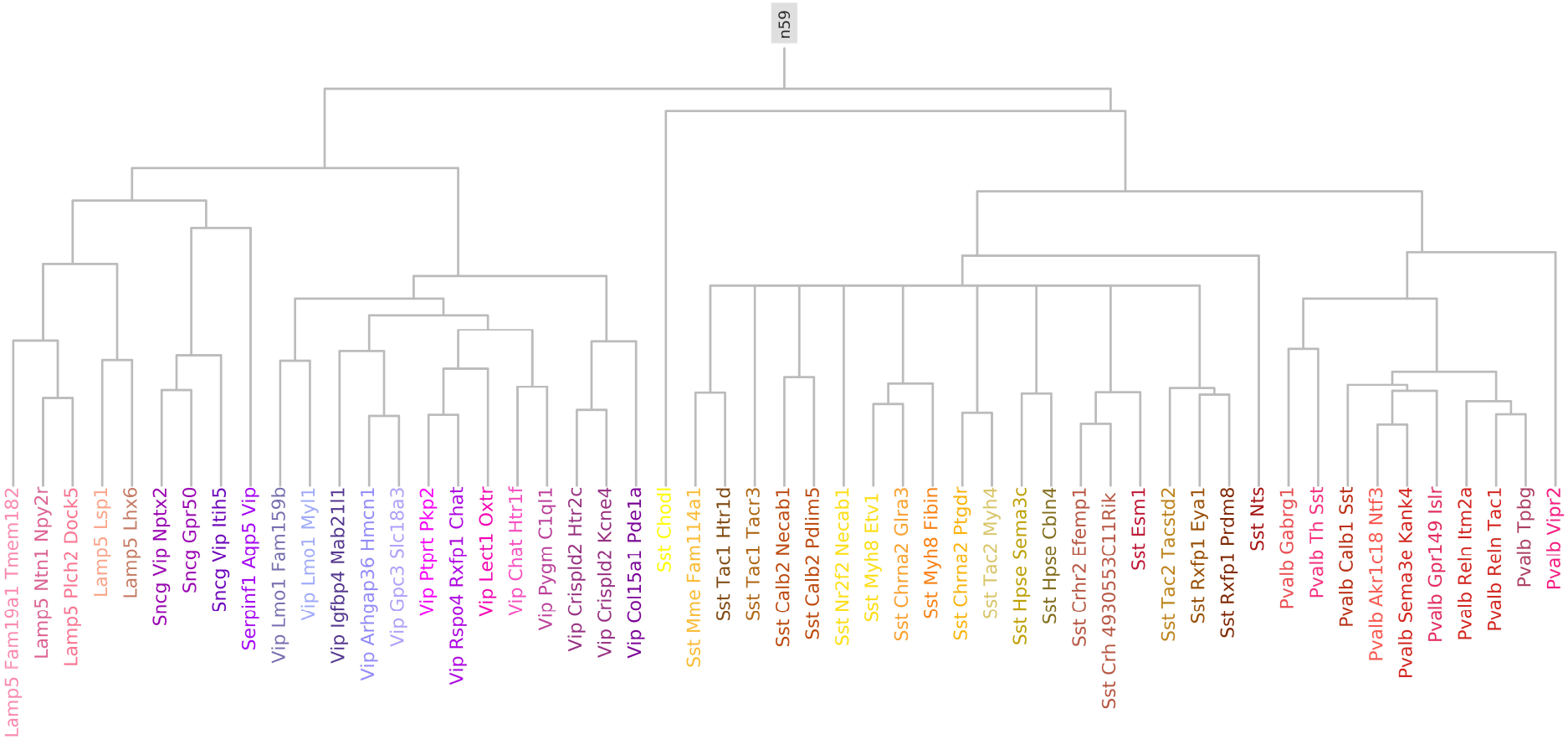
Reference taxonomy for well-represented GABAergic neurons. Cells were mapped to the complete hierarchical classification tree for cortical cells with a marker gene based procedure. Here we show a subset of the full hierarchical tree, which consists of only those leaf nodes that are well-represented (n*≥*10) in the Patch-seq dataset. At the highest resolution, this tree consists of 53 cell type labels. The lowest resolution view consists of a single label (n59) which comprises of all GABAergic cortical neurons.

**Extended Data Figure 2:**
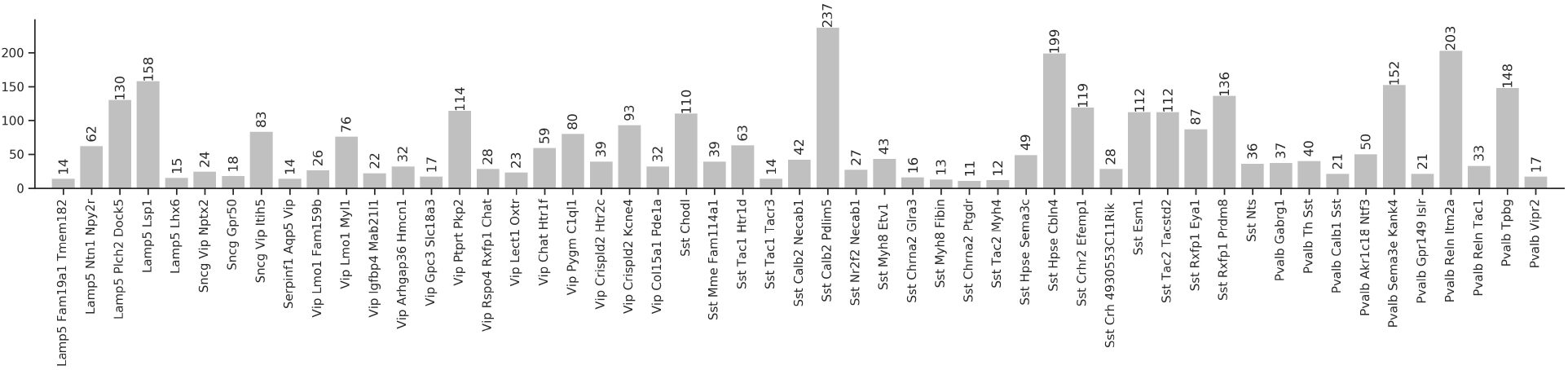
Cell type distribution. The distribution of samples according to the reference hierarchy cell type label assignment. Types with less than 10 samples are not shown.

**Extended Data Figure 3:**
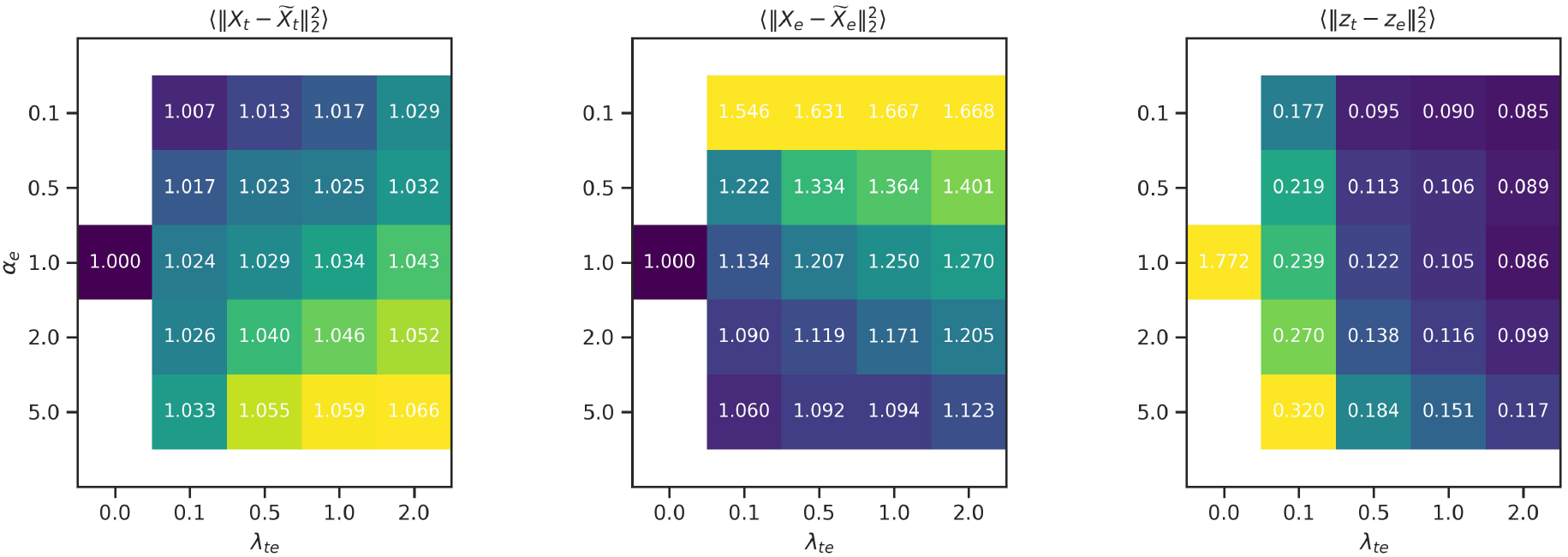
Hyper-parameter search. (Left and center) Reconstruction errors relative to the value over uncoupled networks, and (Right) coupling error over different values for *α*_e_ and *λ*_te_ averages over validation sets. The value for *α*_t_ was set to 1.0 and representation dimensionality was set to 3 for these experiments. As coupling is increased, the reconstruction error increases illustrating the trade-off between coupling and reconstruction accuracy.

**Extended Data Figure 4:**
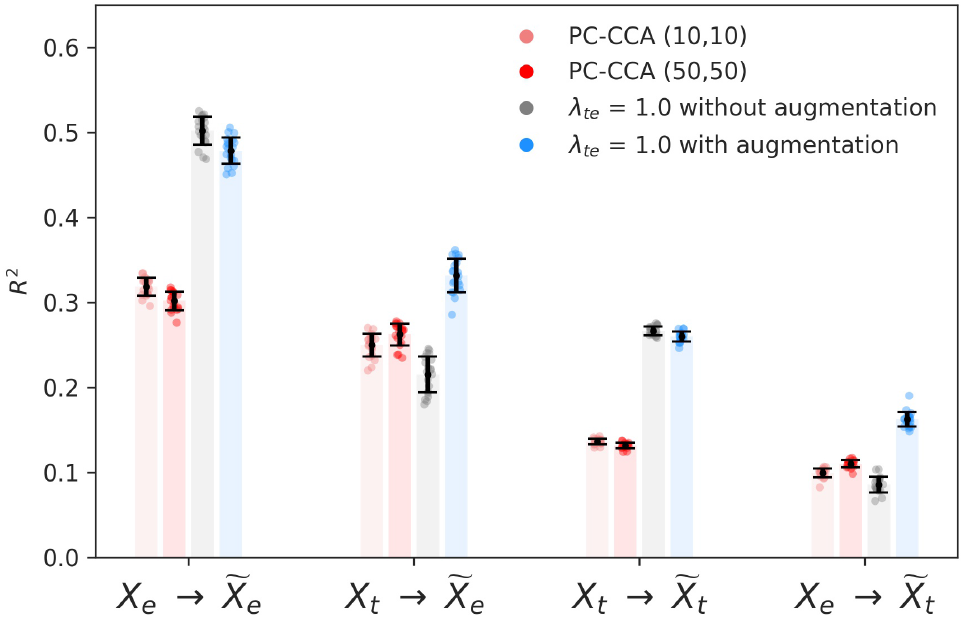
Decoder augmentation improves cross-modal prediction accuracy. We use cross modal representations to augment the input for decoder subnetworks while training. Reconstruction performance as measured by the coefficient of determination (*R*^2^) for linear baselines (PC-CCA), and coupled autoencoders with- and without-augmentation. Error bars show standard deviation over 20 cross validation folds.

**Extended Data Figure 5:**
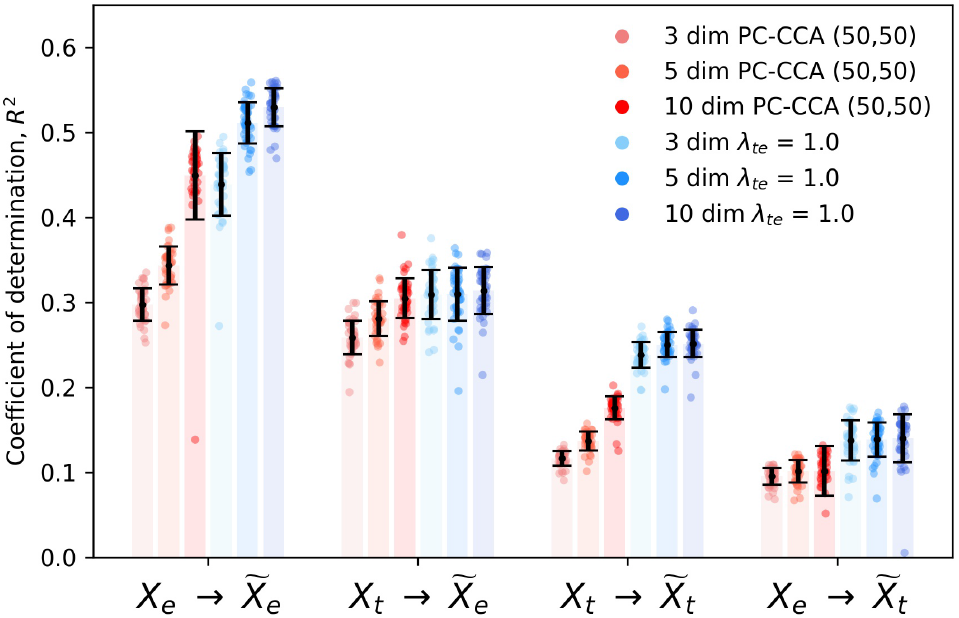
Effect of latent space dimensionality on reconstruction performance. Reconstruction errors for coupled autoencoder and linear baseline for different latent space dimensionality dim *∈* {3, 5, 10}. Coupled autoencoders reconstruct the data more accurately than linear baselines (*p <* 10^*−*4^, two-sided Wilcoxon signed-rank test). The only exception is for 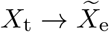 with dimensionality set to 10, where the null hypothesis cannot be rejected. We would like the dimensionality to be as low as possible for downstream tasks such as clustering and classification with limited data, and as high enough for good performance at tasks such as data imputation or cross-modal data prediction.

**Extended Data Figure 6:**
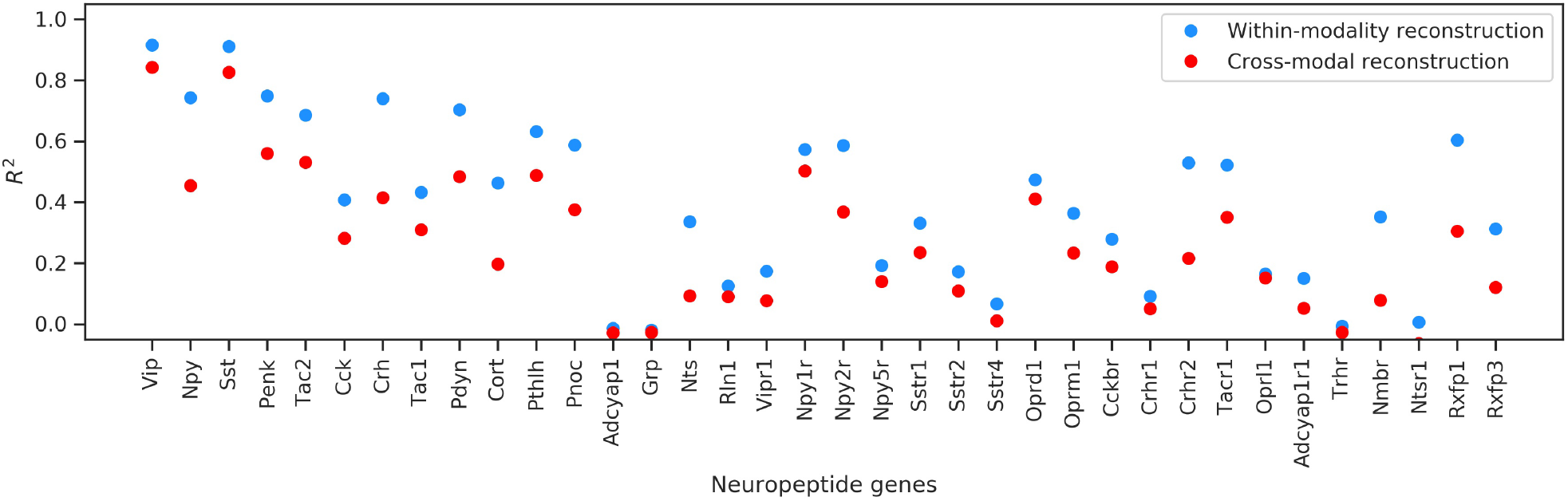
Predicting gene expression from coordinated representations. Within-modality reconstructions for individual genes are decoded from the co-ordinated *λ*_te_ = 1.0 representation *z*_t_ obtained for the transcriptomic data. Cross-modal reconstructions are obtained from the corresponding *z*_e_, which is the representation for the electrophysiological data. The cross-modal reconstructions are comparable to within-modality reconstructions, and a majority of the neuropeptide precursor genes are reconstructed well, as suggested by the high coefficient of determination (*R*^2^) values.

**Extended Data Figure 7:**
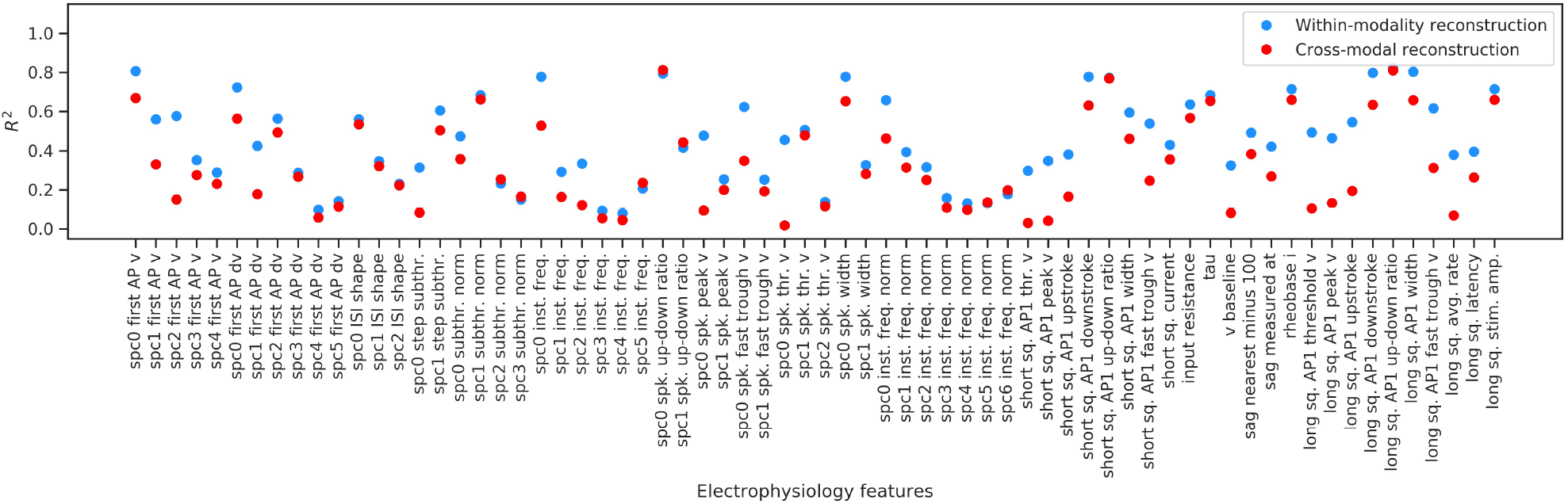
Predicting electrophysiological features based on co-ordinated representations. The within-modality reconstructions for electrophysiological features are decoded from the co-ordinated *λ*_te_ = 1.0 representation *z*_e_ obtained for the electrophysiological data. Cross-modal reconstructions are obtained from the corresponding *z*_t_, which is the representation for the transcriptomic data. Features that are reconstructed well in the within-modality case are analyzed in the context of transcriptomic cell types in the main text.

**Extended Data Figure 8:**
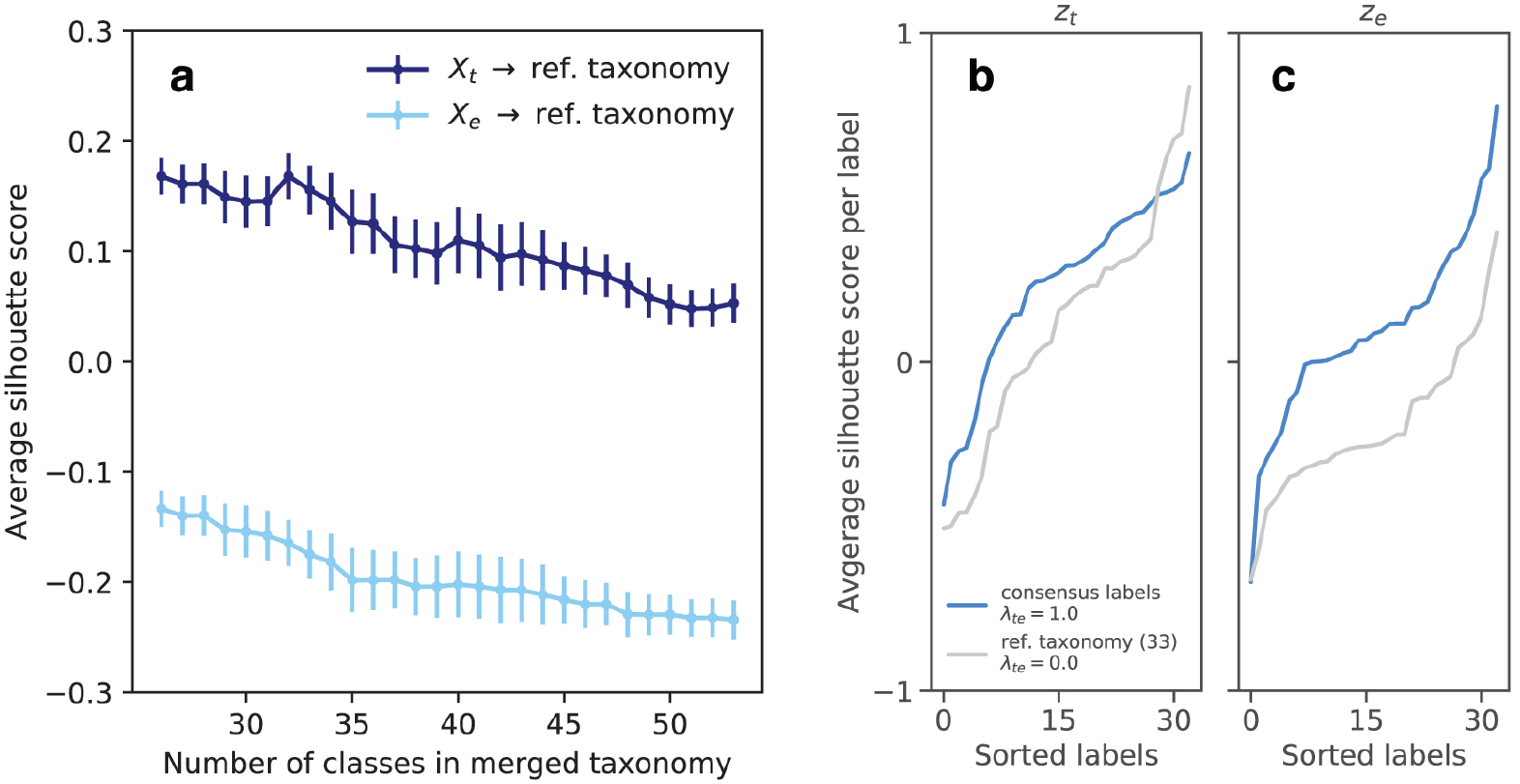
Reference taxonomy labels do not partition the data well. Average silhouette scores for test samples, for successive mergings of the reference taxonomy with uncoupled representations do not indicate any particularly favorable number of clusters. Error bars show mean *±* SD over 5 best initializations of single modality (uncoupled) autoencoders operating on *X*_t_ and *X*_e_. Here the uncoupled representations *z*_t_ and *z*_e_ serve as low dimensional representations of the standalone data. The per-label silhouette score for the 33-merged reference taxonomy labels with uncoupled representations performs worse than consensus cluster labels on both, *z*_t_ (b) and *z*_e_ (c).

**Extended Data Figure 9:**
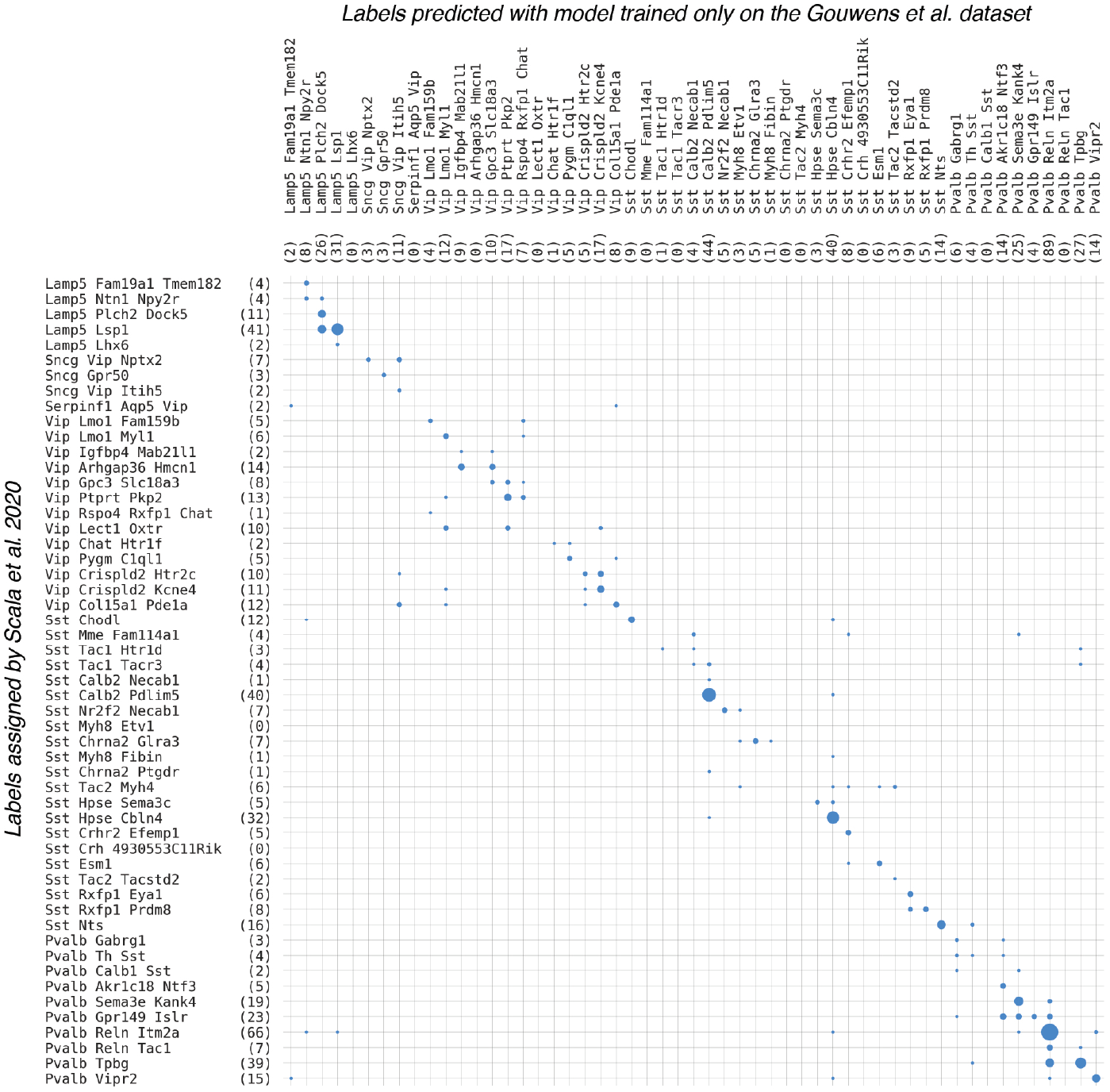
Predicting cell types based on gene expression. The gene expression profiles were used to obtain 3-d representations without additional training. QDA classifiers trained to predict cell types for the Gouwens et al. dataset were used to predict labels for cells in the Scala et al. 2020 dataset, and the resulting contingency matrix is shown. Overall accuracy of label prediction is 66%, with many inaccuracies being accounted for by closely related types.

**Extended Data Figure 10:**
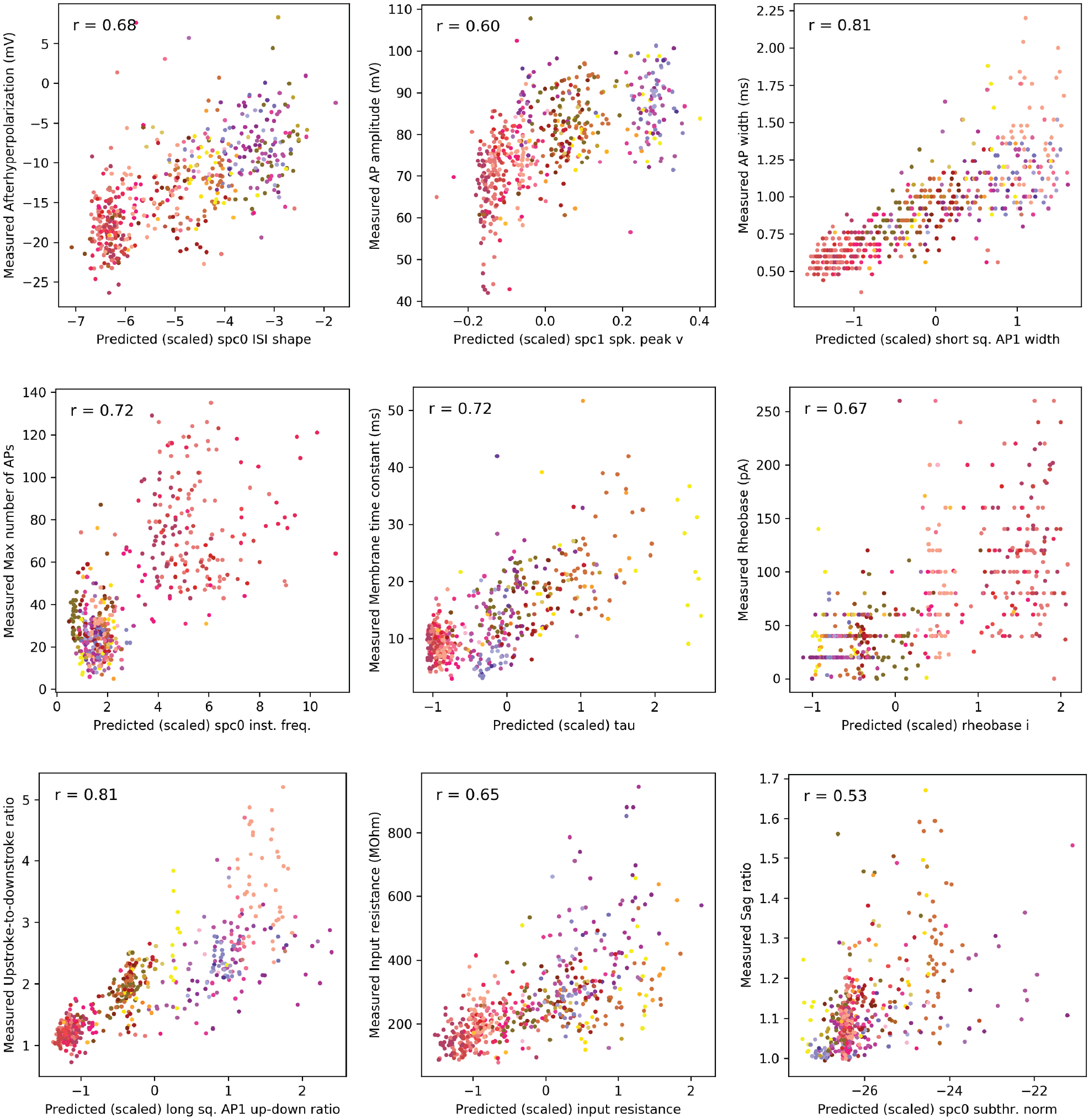
Predicting electrophysiological properties from gene expression. Gene expression profiles for 524 inhibitory neurons in the Scala et al. 2020 dataset were used as input for the coupled autoencoder trained only with the Gouwens et al. dataset. The electrophysiological measurements were not measured the same way in the two datasets; cross-modal setting only allows predictions for electrophysiological features of the Gouwens et al. dataset for cells in the Scala et al. dataset. There is a strong correlation (Pearson’s *r* is shown on each plot) for many related measurements across the datasets. Cells are colored according to the cell type assignments of Scala et al. 2020, who mapped them to the same reference taxonomy that is used throughout this study.

