## Supplementary Information for "Consistent cross-modal identification of cortical neurons with coupled autoencoders"

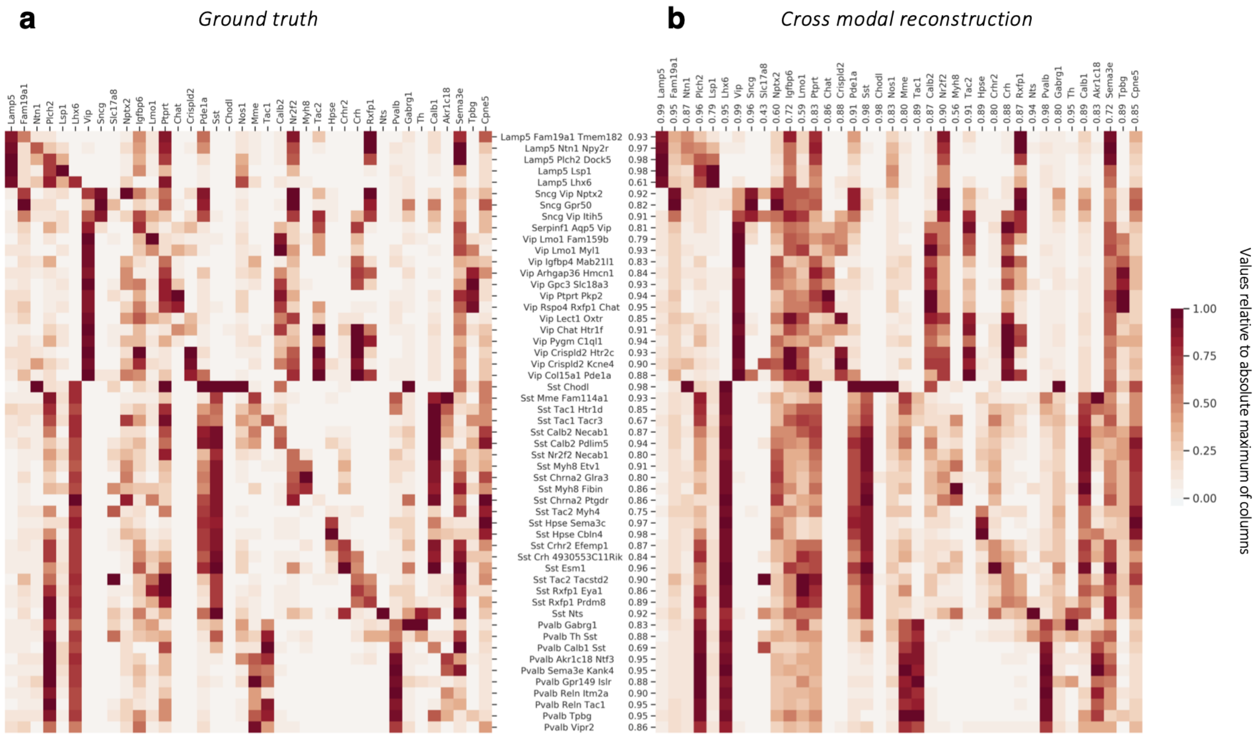


Supplementary Figure 1: **Marker gene expression can be reliably reconstructed from electrophysiological profiles :** (a) Gene expression levels averaged over samples of individual cell types of the reference taxonomy, normalized per gene by the maximum value of each column. (b) Cross-modal reconstructions of these genes using only the electrophysiological features. High row- and column-wise Pearson's $r$ for the reconstructions compared to ground truth indicate that marker genes can be reconstructed with high fidelity starting from electrophysiological features.

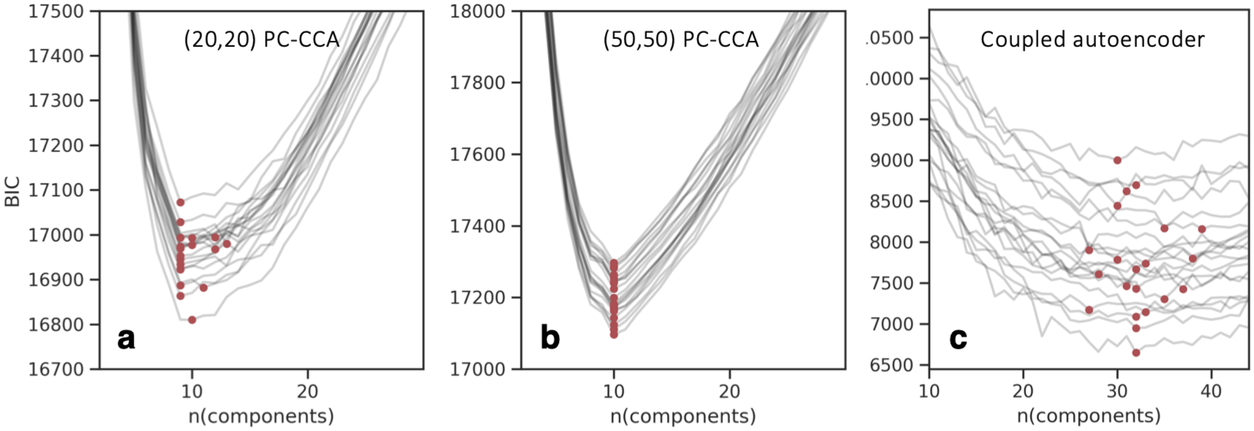


Supplementary Figure 2: **Unsupervised clustering with PC-CCA representation:** As a check, we used the BIC based model selection procedure to determine consistent clusters for the Patch-seq dataset using 3 dimensional PC-CCA representations. A sweep over different number of mixture components was performed, and BIC values are shown for 20 different cross-validation folds for (20,20) PC-CCA in (a) and for (50,50) PC-CCA in (b). The optimal number of clusters is based on the number of components for which the minimum value of BIC is observed for a given representation (red dots). The number of clusters suggested by this procedure is $9.75 \pm 1.22$ for (20,20) PC-CCA and $10 \pm 0$ for (50,50) PC-CCA, in contrast to the $32.19 \pm 3.16$ clusters (mean $\pm$ SD over the different initializations) suggested by the same procedure applied to coupled autoencoder representations, (c). The number of clusters determined by PC-CCA is significantly lower, reinforcing the idea that linear methods fail to identify the variability in cortical GABAergic neurons below the subclass level.

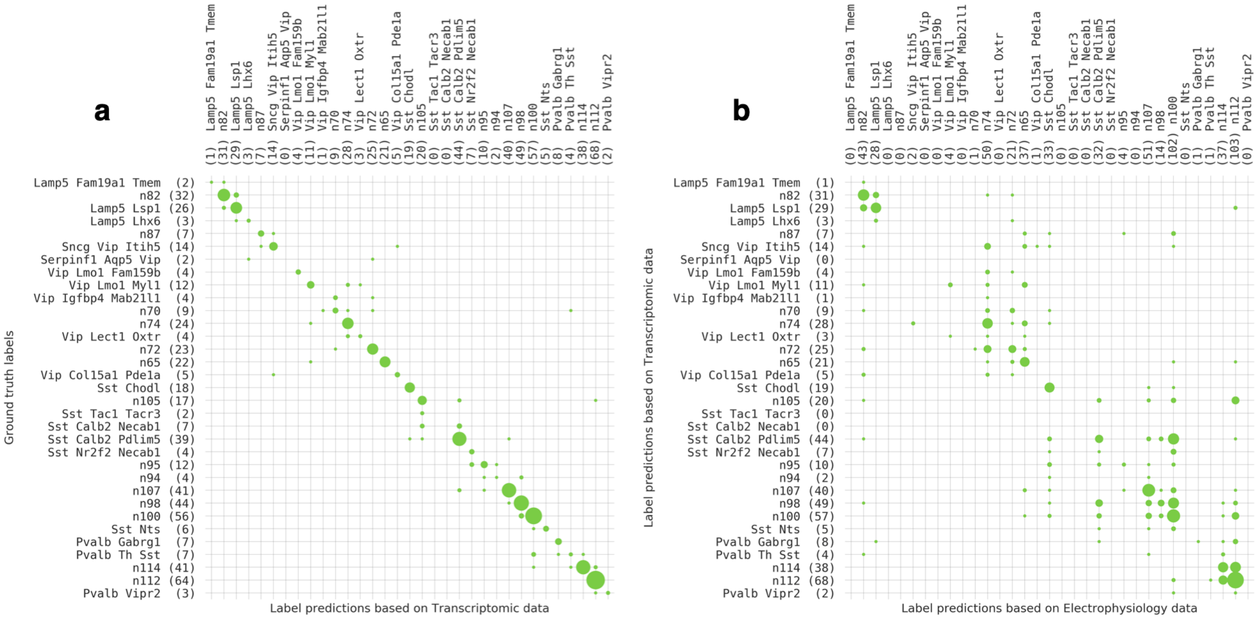


Supplementary Figure 3: **Evaluating a supervised approach for consensus of labels based on the reference taxonomy:** (a) Representative contingency matrix with test samples, comparing 33-way merged reference taxonomy labels with classifier label predictions using only the transcriptomic data shows that an independent, 3-dimensional autoencoder representation can be used effectively for classification. The 33-way merged reference taxonomy labels are obtained by slicing the hierarchy in Figure 1 with a single horizontal cut. Labels such as n74, n70 etc. are non-leaf internal nodes of the hierarchical taxonomy. (b) Comparing the labels assigned by a classifier trained on the transcriptomic representation with one independently trained on the electrophysiology representation is noisy, with many labels that are not assigned at all, suggesting that electrophysiology data by itself is not characterized well by the reference transcriptomic taxonomy.

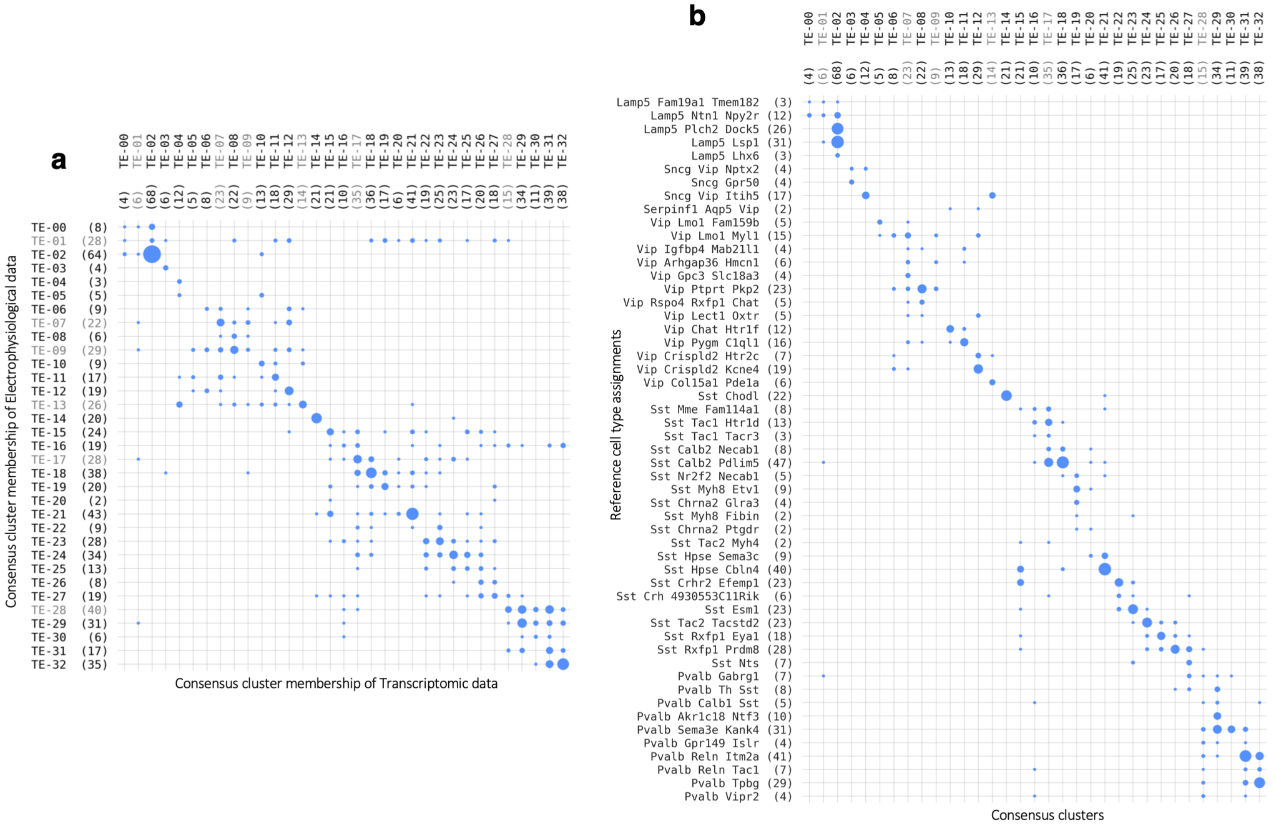


Supplementary Figure 4: **Consensus clusters for test cells** Contingency matrices to evaluate consensus between the modalities (a) and to compare with reference taxonomy (b) for the subset (∼20%) of cells in Figure 3 that were not part of autoencoder training, nor part of training the Gaussian mixture models.

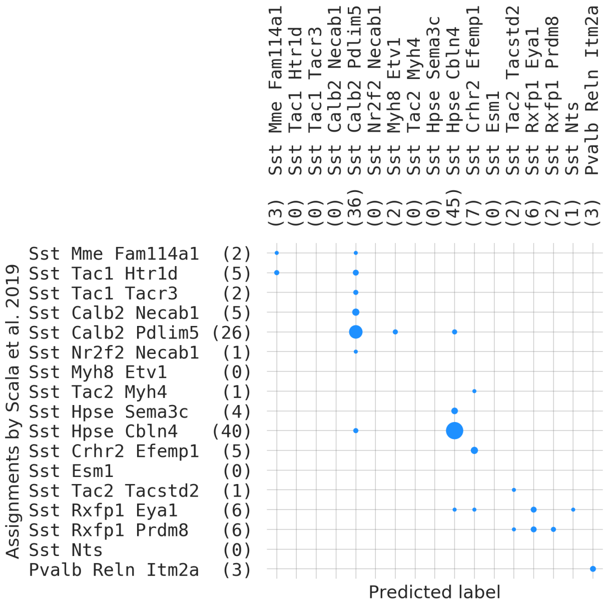


Supplementary Figure 5: **Predicting cell types based on gene expression:** Gene expression measurements for 107 cells in the Scala et al. 2019 dataset are used as input to one of the coupled autoencoders trained on the Gouwens et al. dataset. A QDA cell type classifier is trained on the aligned representation of the Gouwens et al. dataset, and is used to predict the type for the cells in the Scala et al. 2019 dataset. The labels assigned by Scala et al. are used as ground truth to construct the contingency matrix. The overall accuracy of label prediction is $> 70\%$ , with many inaccuracies being accounted for by closely related types (the cell types in the plot are ordered according to the reference hierarchical taxonomy used in this study.)

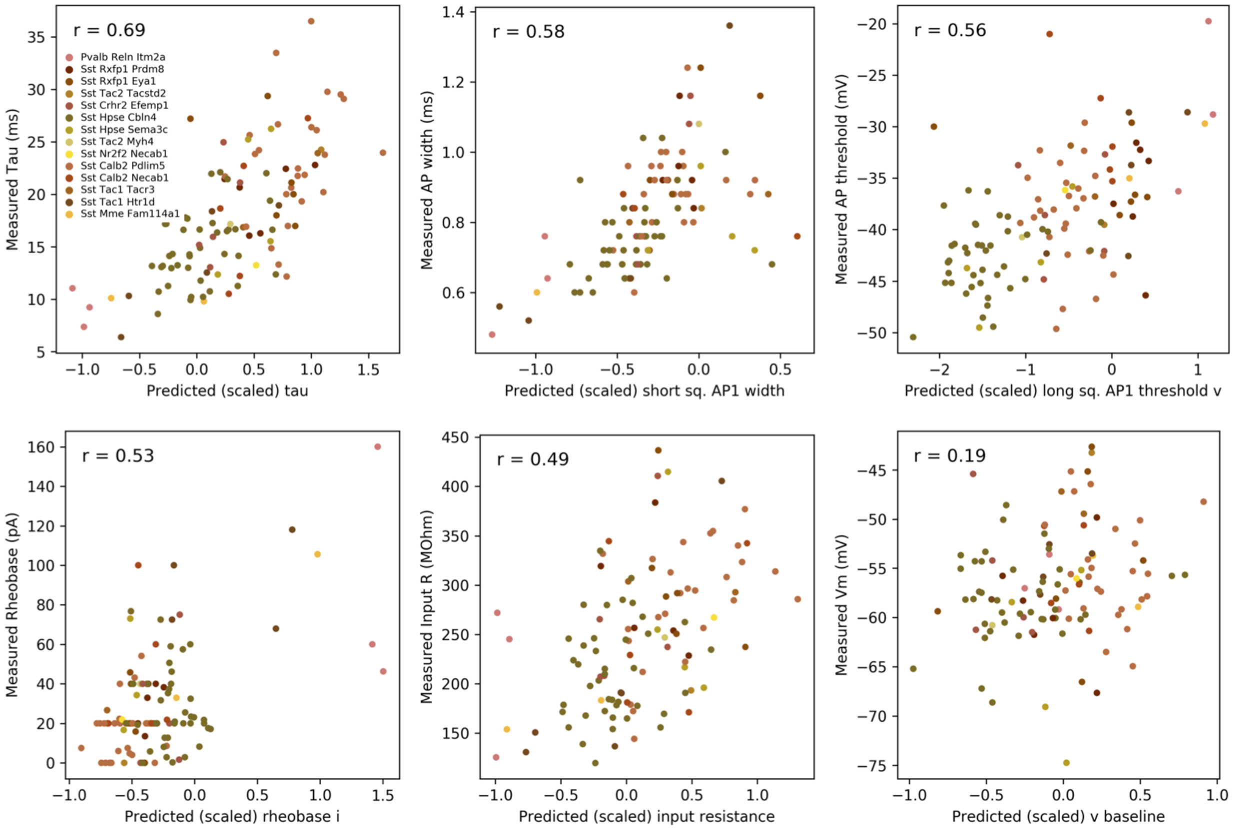


Supplementary Figure 6: **Predicting electrophysiological properties from gene expression:** Gene expression profiles for 107 cells in the Scala et al. dataset were used as input for the coupled autoencoder trained only with the Gouwens et al. dataset. The electrophysiological measurements were not measured the same way in the two datasets; cross-modal setting only allows predictions for electrophysiological features of the Gouwens et al. dataset for cells in the Scala et al. dataset. There is a strong correlation (Pearson's $r$ is shown on each plot) for many related measurements across the datasets. Cells are colored according to the cell type assignments of Scala et al. 2019, who mapped them to the same reference taxonomy that is used throughout this study.

### Probabilistic setting

The deterministic view in Eq. 4 (Methods) is equivalent to maximizing log-likelihood of a discriminative probabilistic model for i.i.d. observations:

$$\begin{aligned}\sum_{s\in S}\log p(x_{st},x_{se},\hat{z}_{st}|\hat{z}_{se}) &= \sum_{s\in S}\log p(x_{st}|\hat{z}_{st},\hat{z}_{se})+\log p(x_{se}|\hat{z}_{se})+\log p(\hat{z}_{st}|\hat{z}_{se})\\ &= \sum_{s\in S}\log p(x_{st}|\hat{z}_{st})+\log p(x_{se}|\hat{z}_{se})+\log p(\hat{z}_{st}|\hat{z}_{se}), \qquad (1)\end{aligned}$$

where we assume that $x_{se}$ is independent of $x_{st}$ and $\hat{z}_{st}$ given $\hat{z}_{se}$, and $x_{st}$ is independent of $\hat{z}_{se}$ given $\hat{z}_{st}$. Modeling the conditional probabilities with Gaussian distributions, $x_{st}|\hat{z}_{st} \sim \mathcal{N}(\widetilde{x}_{st}, \sigma_{\mathrm{t}}^2 I)$, $x_{se}|\hat{z}_{se} \sim \mathcal{N}(\widetilde{x}_{se}, \sigma_{\mathrm{e}}^2 I)$, and $\hat{z}_{st}|\hat{z}_{se} \sim \mathcal{N}(\hat{z}_{se}, \lambda^{-1} I)$, we can write Eq. 1 as

$$\sum_{s\in S} \log p(x_{st}, x_{se}, \hat{z}_{st}|\hat{z}_{se}) = \frac{-1}{2} \sum_{s\in S} \sigma_{\mathrm{t}}^{-2} ||x_{st} - \widetilde{x}_{st}||_2^2 + \sigma_{\mathrm{e}}^{-2} ||x_{se} - \widetilde{x}_{se}||_2^2 + \lambda \|\hat{z}_{st} - \hat{z}_{se}\|_2^2 + \text{const.} \quad (2)$$

Comparing the loss function in Eq. 2 with Eq. 4 (Methods), we see that $\alpha_{\mathrm{t}} \sim \sigma_{\mathrm{t}}^{-2}$ and $\alpha_{\mathrm{e}} \sim \sigma_{\mathrm{e}}^{-2}$ are related to the noise in the measurements, and $\lambda_{\mathrm{te}} \sim \lambda$ denotes the precision in cross-modal latent variable estimation.

The two modalities t and e are interchangeable in Eq. 1, and Figure 1b suggests that the individual cell types may be well-approximated by hyperellipsoids. Therefore, fitting a Gaussian mixture model to the encodings provides an efficient prior distribution for $p(\hat{z}_{se})$ (or $p(\hat{z}_{st})$), and produces a generative model for multimodal datasets.

#### Overview of alignment methods

**Distribution matching methods**: Approaches to align multimodal datasets either in the data domain or in a low dimensional latent space using generative adversarial losses, optimal transport, maximum mean discrepancy loss etc. have been employed for single cell data .[1–4] Nevertheless, by virtue of a lack of pairing, the best they can achieve is a matching at the distribution level, which is not enough for principled alignment of drastically different observation modalities. For instance, unless the transcriptomic and electrophysiological datasets are matched across all observed and unobserved axes of variability (e.g., layer distribution, cell type distribution, region distribution, age/sex distribution of animals), these methods are not applicable to the fine-scale correspondence problem studied in this manuscript.

When applied to paired datasets, these methods may superficially appear to perform well. However, they cannot resolve the symmetries in the distribution. (e.g., symmetries in the individual components of a generalized mixture model). In the presence of a distributional symmetry, these methods cannot guarantee the correct alignment (e.g., $a \to a'$, $b \to b'$): they are as likely to produce swapped mappings (e.g., $a \to b'$, $b \to a'$), by virtue of not using the pairing information.

Lastly, adversarial training owes most of its success to image data. For unstructured datasets (i.e., a list of features), the training process can be unstable, resulting in the practice of reporting the best run out of multiple runs in this field. In contrast, we have not observed such catastrophic training failures with our *coupled autoencoder* method, across thousands of runs.

While it is possible to introduce a pairing loss term to the GAN framework,[1] our approach completely removes the dependence on adversarial training and can also work with partially paired datasets. Thus, our method has three main advantages over such a distribution matching based approach: (i) it naturally produces a latent space representation, which is crucial for many downstream tasks including clustering and visualization, (ii) it avoids training stability issues by not using an adversarial loss term, (iii) it does not impose alignment penalties at the distribution level and instead only relies on self-reconstruction accuracy for unpaired cells, and self/cross reconstruction accuracy and the alignment accuracy for paired cells, thus avoiding forced alignment.

**Methods based on parametric generative models**: While these approaches can efficiently infer underlying factors of variability and offer enhanced interpretability,[5] they require detailed statistical characterizations of the experimental protocol. Such models do not exist for electrophysiology, morphology, connectivity, etc. Therefore, parametric modeling is typically applied only to genetic datasets, mostly for batch correction purposes.

**CCA-based methods**: This family includes CCA, deepCCA, and its derivatives,[6],[7–9] as well as our method. Indeed, our method addresses two computational issues of deepCCA: (i) deepCCA infers explicit transformation matrices in addition to the parameters of the encoder/decoder neural networks. Our method does not depend on such extraneous matrices, thereby decreasing the number of parameters and providing a more intuitive approach. (ii) Our method does not solve the standard eigenvalue problem and instead computes only the minimum singular value. This results in significant computational and performance improvements for small batch sizes and/or large embedding dimensionalities.[10] Importantly, similar earlier attempts[11] failed to produce non-collapsing latent space representations. Finally, we found that augmenting the objective function with cross-modal reconstruction loss significantly improves the alignment accuracy (Extended Data Figure 4).

[2] Pinar Demetci, Rebecca Santorella, Bjorn Sandstede, William Stafford Noble, and Ritambhara Singh. Gromov-wasserstein optimal transport to align single-cell multi-omics data. *BioRxiv*,

2020.

[3] Matthew Amodio, David Van Dijk, Krishnan Srinivasan, William S Chen, Hussein Mohsen, Kevin R Moon, Allison Campbell, Yujiao Zhao, Xiaomei Wang, Manjunatha Venkataswamy, et al. Exploring single-cell data with deep multitasking neural networks. *Nature methods*, pages 1–7, 2019.

[4] Jay S Stanley III, Scott Gigante, Guy Wolf, and Smita Krishnaswamy. Harmonic alignment. In *Proceedings of the 2020 SIAM International Conference on Data Mining*, pages 316–324. SIAM, 2020.

[5] Romain Lopez, Jeffrey Regier, Michael B Cole, Michael I Jordan, and Nir Yosef. Deep generative modeling for single-cell transcriptomics. *Nature methods*, 15(12):1053–1058, 2018.

[6] Trevor Hastie, Robert Tibshirani, and Jerome Friedman. *The elements of statistical learning: data mining, inference, and prediction*. Springer Science & Business Media, 2009.

[7] Francis R Bach and Michael I Jordan. A probabilistic interpretation of canonical correlation analysis. 2005.

[8] Galen Andrew, Raman Arora, Jeff Bilmes, and Karen Livescu. Deep canonical correlation analysis. In *International conference on machine learning*, pages 1247–1255, 2013.

[9] Gregory Gundersen, Bianca Dumitrascu, Jordan T Ash, and Barbara E Engelhardt. End-to-end training of deep probabilistic cca on paired biomedical observations. In *Uncertainty in artificial intelligence*, 2019.

[10] Rohan Gala, Nathan Gouwens, Zizhen Yao, Agata Budzillo, Osnat Penn, Bosiljka Tasic, Gabe Murphy, Hongkui Zeng, and Uygar Sümbül. A coupled autoencoder approach for multi-modal analysis of cell types. In *Advances in Neural Information Processing Systems*, pages 9263–9272, 2019.

[11] Fangxiang Feng, Xiaojie Wang, and Ruifan Li. Cross-modal retrieval with correspondence autoencoder. In *Proceedings of the 22nd ACM international conference on Multimedia*, pages 7–16, 2014.
